# High-throughput Evaluation of Epilepsy-associated *KCNQ2* Variants Reveals Functional and Pharmacological Heterogeneity

**DOI:** 10.1101/2021.10.18.464842

**Authors:** Carlos G. Vanoye, Reshma R. Desai, Zhigang Ji, Sneha Adusumilli, Nirvani Jairam, Nora Ghabra, Nishtha Joshi, Eryn Fitch, Katherine Helbig, Dianalee McKnight, Amanda Lindy, Fanggeng Zou, Ingo Helbig, Edward Cooper, Alfred L. George

## Abstract

Hundreds of *KCNQ2* variants have been identified by genetic testing of children with early onset epilepsy and/or developmental disability. Voltage-clamp recording from heterologous cells has proved useful for establishing deleterious functional effects of *KCNQ2* variants, but procedures adapting these assays for standardized, higher throughput data collection and reporting are lacking. In this study, we employed automated patch clamp recording to assess in parallel the functional and pharmacological properties of 79 missense and 2 in-frame deletion variants of *KCNQ2*. Among the variants we studied were a training set of 18 pathogenic variants previously studied by voltage-clamp recording, 24 mostly rare population variants, and 39 disease-associated variants with unclear functional effects. Variant KCNQ2 subunits were transiently expressed in a cell line stably expressing KCNQ3 to reconstitute the physiologically relevant channel complex. Variants with severe loss-of-function were also co-expressed 1:1 with WT KCNQ2 in the KCNQ3 cell line to mimic the heterozygous genotype and assess dominant-negative behavior. In total, we analyzed electrophysiological data recorded from 9,480 cells. The functional properties of WT KCNQ2/KCNQ3 channels and pharmacological responses to known blockers and activators determined by automated patch clamp recording were highly concordant with previous findings. Similarly, functional properties of 18 known pathogenic variants largely matched previously published results and the validated automated patch clamp assay. Many of the 39 previously unstudied disease-associated KCNQ2 variants exhibited prominent loss-of-function and dominant-negative effects, providing strong evidence in support of pathogenicity. All variants, exhibit response to retigabine (10 µM), although there were differences in maximal responses. Variants within the ion selectivity filter exhibited the weakest responses whereas retigabine had the strongest effect on gain-of-function variants in the voltage-sensor domain. Our study established a high throughput method to detect deleterious functional consequences of KCNQ2 variants. We demonstrated that dominant-negative loss-of-function is a common mechanism associated with missense KCNQ2 variants but this does not occur with rare population variation in this gene. Importantly, we observed genotype-dependent differences in the response of KCNQ2 variants to retigabine.

## INTRODUCTION

Pathogenic variants in voltage-gated sodium and potassium channels genes account for a large fraction of individuals diagnosed with early life epilepsy and related developmental and epileptic encephalopathies (DEEs).^1-3,4^ Recent investigations of disease-associated variants have yielded important discoveries about the molecular mechanisms responsible for early life epilepsy and inspired a framework for individualized treatment known as precision medicine,^5,6^ However, increased use of genetic testing has resulted in the ongoing discovery of a greater number of candidate pathogenic variants than can be accommodated by standard investigative approaches.

An important cause of monogenic epilepsy, *KCNQ2*, encodes a voltage-gated potassium subunit (KCNQ2 or K_V_7.2) that forms tetrameric channels by itself, or co-assembles with the related KCNQ3/K_V_7.3 subunit to generate an important, slowly-activating, non-inactivating muscarinic acetylcholine receptor modulated current (M-current) in central and peripheral neurons.^7-10^ *KCNQ2* related disorders are highly penetrant, but exhibit phenotypic and allelic heterogeneity that makes laboratory and clinical variant interpretation challenging. Benign familial neonatal epilepsy (BFNE) is dominantly-transmitted and caused in about 70% of pedigrees by *KCNQ2* stop-gain variants.^11-13^ Notably, *KCNQ2* and certain *KCNQ3* variants cause BFNE that is phenotypically indistinguishable. By contrast, other *KCNQ2* variants, usually missense, or, more rarely, small in-frame deletions arising *de novo*, have been identified in persons with a spectrum of more disabling phenotypes including neonatal onset seizures with subsequent moderate to severe global disability, and later-onset disability with and without seizures.^14-17^ Elucidation of genotype-phenotype relationships along the *KCNQ2* DEE spectrum has relied on identification and multidisciplinary study of a handful of recurrent *KCNQ2* variants. These studies have revealed correlation between phenotypes and distinctive functional profiles, ranging from strong loss-of-function^18,19^ to strong gain-of-function.^14-16^ Although these foundational studies provided a framework for *KCNQ2* variant genotype-phenotype correlation, several hundred disease-associated *KCNQ2* variants have unknown functional consequences. The lack of functional evidence regarding these variants limits assignment of pathogenicity, leaves individual families’ diagnostic journey in limbo, and may be a hurdle to enrollment into precision medicine trials.

*In vitro* voltage-clamp studies of heterologously expressed recombinant human K_V_7.2 channels have been informative regarding the functional consequences of disease-associated variants.^12,18-25^ Although valuable, these studies have been confounded by varied experimental strategies and expression systems employed by multiple independent laboratories. This experimental heterogeneity makes direct comparison of findings for variants studied in different laboratories and by different methods challenging. Furthermore, targeted therapy of *KCNQ2*-related epilepsy with an approved small molecule M-current activator (retigabine/ezogabine) has been demonstrated,^11^ but whether all disease-associated variants will respond equally to this drug has not been determined.

In this study, we developed a pipeline for profiling key functional consequences of *KCNQ2* variants with a rigorously validated strategy using automated patch clamp recording. This approach enabled us to compare the functional properties of disease-associated variants with rare population variants, and to systematically investigate the pharmacological response of individual variants to retigabine. Our findings greatly expand knowledge regarding the molecular basis of *KCNQ2*-related epilepsy including functional comparisons among variants associated with the spectrum of clinical severity, and offer new information about the heterogeneity of retigabine responsiveness that will contribute to its emerging use for precise therapy of this monogenic epilepsy.

## MATERIALS AND METHODS

### Variant identification and prioritization for study

To enable validation and deployment of the automated patch recording pipeline, we prioritized 81 variants from de-identified data provided by a clinical testing laboratory (GeneDX, Gaithersburg MD), public databases (Human Gene Mutation Database [HGMD], gnomAD, RIKEE), and de-identified clinical research databases maintained at Baylor College of Medicine, Northwestern University, and the Children’s Hospital of Philadelphia. All clinical data were collected after informed consent under approved IRB protocols and were de-identified prior to consideration for this study. Variants were sorted into categories based on availability of phenotypes, recurrence, location in pathogenic hotspots,^11,12,25^ and previously published functional characterization. Our study included variants with well-described phenotypes (including BFNE, neonatal-onset DEE, and later-onset DEE with variable seizure severity) and previous comprehensive evaluations by voltage-clamp electrophysiology. Additional sets of variants with incompletely understood genotype-phenotype relationships, were also selected. These variants included several highly recurrent variants not yet described in published functional studies, rare variants for which clinical phenotyping was less extensive, and variants where earlier voltage-clamp study was fragmentary, or merited re-testing due to atypical or conflicting results. Based on gnomAD, we also selected 23 rare variants (minor allele frequency ranging from 0.000008 to 0.003) and a single *KCNQ2* common variant (N780T; **Table S1**). All variants were annotated based on the NCBI reference sequence NM_172104.

### Cell culture

Chinese hamster ovary cells (CHO-K1; CRL 9618, American Type Culture Collection, Manassas VA, USA) were grown in F-12 nutrient mixture medium (GIBCO/Invitrogen, San Diego, CA, USA) supplemented with 10% fetal bovine serum (ATLANTA Biologicals, Norcross, GA, USA), penicillin (50 units·ml^-1^), and streptomycin (50 μg·ml^-1^) at 37°C in 5% CO_2_.

A cell line (CHO-Q3) stably expressing human wild type (WT) KCNQ3 (Q3, GenBank accession number NM_004519) was generated using Flp-In™-CHO cells according to the supplier’s protocol (Thermo Fisher Scientific, Waltham, USA). CHO-Q3 stable cells were maintained under dual selection with Zeocin (100 μg/ml) and hygromycin B (600 μg/ml) in F-12 nutrient mixture medium (GIBCO/Invitrogen) supplemented with 10% fetal bovine serum (ATLANTA Biologicals), penicillin (50 units·ml^-1^), and streptomycin (50 μg·ml^-1^) at 37°C in 5% CO_2_. Q3-expressing clones were screened by electroporating WT KCNQ2 and measuring M-current density by automated patch clamp recording (see below). One clone (L1) exhibiting the highest percentage of cells expressing M-current (defined as XE-991-sensitive current) was selected for experiments. The cell line was only used between passage number 4 and 15.

Unless otherwise stated, all tissue culture media were obtained from Life Technologies, Inc. (Grand Island, NY, USA).

### Plasmids and transfection

A cDNA encoding the complete open reading frame of KCNQ3 was cloned into pcDNA5/FRT for use in generating the CHO-Q3 stable cell line. Other vectors were designed to enable co-expression of untagged channel subunits with fluorescent proteins as a means for tracking successful cell transfection. A cDNA with the complete coding region of human KCNQ2 (GenBank accession number NM_172108, which is an abundant splice isoform in human brain^26^ was engineered in the mammalian expression vector pIRES2-EGFP (BD Biosciences-Clontech, Mountain View, CA, USA). The EGFP-expressing vector was used for transfection of WT or variant KCNQ2 into CHO-Q3 cells (homozygous state). For studies designed to investigate KCNQ2 variants in the heterozygous state, CHO-Q3 cells were co-transfected with KCNQ2 variant in the EGFP-expressing vector and KCNQ2-WT in a modified vector where EGFP was substituted with the cyan-excitable orange-red fluorescent protein CyOFP1.^27^

Three KCNQ2 variants (A193D, R201C, P335L) yielded too few positively transfected cells for automated patch clamp recording when expressed as homozygous channels in CHO-Q3 cells. We overcame this limitation by transiently co-transfecting CHO-K1 cells with each of these variants along with KCNQ3, which was expressed using a modified pIRES2-vector co-expressing the monomeric red fluorescent protein mScarlet.^28^ These variants were successfully analyzed using CHO-Q3 cells when expressed as heterozygous channels.

### Site-directed mutagenesis

KCNQ2 variants were introduced into the WT coding sequence using Q5 High Fidelity DNA polymerase (New England Biolabs, Ipswich, MA, USA). Mutagenic primers for each variant were designed using custom software (Q5 Designer; source code available upon request). Mutagenic primer sequences are provided in **Table S2**). Variant KCNQ2 plasmid clones were screened as previously described for the related potassium channel KCNQ1.^28^ For each variant plasmid, the complete KCNQ2 coding region, the IRES element and fluorescent protein cDNA were sequenced in their entirety by Sanger sequencing (Eurofins Genomics, Louisville, KY) and analyzed using a custom multiple sequence alignment tool (Multiple Sequence Iterative Comparator [MuSIC], source code available upon request). Endotoxin-free plasmid DNA was purified from clones with correct sequences (Nucleobond Xtra Maxi EF, Macherey-Nagel Inc.), and re-suspended in endotoxin-free water.

### Electroporation

Plasmids encoding WT or variant KCNQ2 channels were transiently transfected by electroporation using the Maxcyte STX system (MaxCyte Inc., Gaithersburg, MD, USA).^29^ CHO-Q3 or CHO-K1 cells grown to 70-80% confluence were harvested using 0.25% trypsin. A 500 µl aliquot of cell suspension was then used to determine cell number and viability using an automated cell counter (ViCell, Beckman Coulter). Remaining cells were collected by gentle centrifugation (160 × g, 4 minutes), washed with 5 ml electroporation buffer (EBR100, MaxCyte Inc.), and re-suspended in electroporation buffer at a density of 10^8^ viable cells/ml. Each electroporation was performed using 100 µl of cell suspension.

CHO-Q3 cells were electroporated with 15 µg of WT or variant KCNQ2 cDNA for homozygous channel studies or with 10 µg of each variant and WT KCNQ2 cDNAs for experiments testing the heterozygous state. For the 3 variants tested by co-expression with KCNQ3 in CHO-K1 cells, 15 μg of variant or WT KCNQ2 cDNA and 35 μg of KCNQ3 cDNA were used per electroporation. The DNA-cell suspension mix was transferred to an OC-100 processing assembly (MaxCyte Inc.) and electroporated using the preset CHO-PE protocol. Immediately after electroporation, 10 µl of DNase I (Sigma-Aldrich, St. Louis, MO, USA) was added to the DNA-cell suspension and the entire mixture was transferred to a 35 mm tissue culture dish for a 30 min incubation at 37°C in 5% CO_2_. Following incubation, cells were gently re-suspended in culture media, transferred to a T75 tissue culture flask and grown for 48 hours at 37°C in 5% CO_2._Following incubation, cells were harvested, counted, transfection efficiency determined by flow cytometry (see below), and then frozen in 1 ml aliquots at 1.5 × 10^6^ viable cells/ml in liquid N_2_ until used in experiments.

### Flow Cytometry

Transfection efficiency was evaluated by flow cytometry (CytoFLEX, Beckman Coutler) using a 488 nm laser. Forward scatter (FSC), side scatter (SSC), green fluorescence (FITC, EGFP), orange fluorescence (PEA, CyOFP) and red fluorescence (PE, mScarlet) were recorded. FSC and SSC were used to gate single viable cells and to eliminate doublets, dead cells and debris. Ten thousand events were recorded for each sample. Non-electroporated cells were assayed as control for all parameters to set the gates for each experiment, the percentage of fluorescent cells was determined from the gated population. Compensation matrixes were generated using CHO-Q3 or CHO-K1 cells expressing only single fluorescent markers and applied to the co-transfected cells to account for spectrum overlap. The percentage of co-transfected cells was determined from plots of FITC *vs* PEA or FITC *vs* PE fluorescence intensity.

### Cell preparation for automated electrophysiology

Electroporated cells were thawed the day before experiments, plated and incubated for 10 hours at 37°C in 5% CO_2_. The cells were then grown overnight at 28°C in 5% CO_2_ to increase channel expression at the plasma membrane. Prior to experiments, cells were passaged using 5% trypsin in cell culture media. Cell aliquots (500 µl) were used to determine cell number and viability by automated cell counting, and transfection efficiency by flow cytometry. Cells were then diluted to 200,000 (single-hole chips) or 600,000 (4-hole chips) cells/ml with external solution (see below) and allowed to recover 60 minutes at 15°C while shaking on a rotating platform at 200 rpm.

### Automated patch clamp recording

Automated patch clamp recording was performed using the Syncropatch 768 PE platform (Nanion Technologies, Munich, Germany) as previously described.^28,29^ Experiments were performed blinded to variant phenotype or source through to the completion of data analysis. Single-hole, 384-well recording chips with medium resistance (2-4 MΩ) were used for most experiments. Four-hole, 384-well recording chips with medium resistance were used to investigate TEA dose-response relationships. The external solution contained (in mM): NaCl 140, KCl 4, CaCl_2_ 2, MgCl_2_ 1, HEPES 10, glucose 5, with the final pH adjusted to 7.4 with NaOH. The internal solution contained (in mM): KF 60, KCl 50, NaCl 10, HEPES 10, EGTA 10, ATP-Mg 5, with the final pH adjusted to 7.2 with KOH.

Pulse generation and data collection were carried out with PatchController384 V.1.3.0 software (Nanion Technologies, Munich, Germany). Whole-cell currents recorded at room temperature in the whole-cell configuration were filtered at 3 kHz and acquired at 10 kHz. The access resistance and apparent membrane capacitance were estimated using built-in protocols. Access resistance compensation was set to 80%. Stringent criteria were used to include individual recordings for final data analysis. We required seal resistance to be ≥0.5 gigaohm except for recordings performed after application of XE-991 where we used a threshold of ≥0.3 gigaohm and recordings done with 4-hole chips in which the criterion for inclusion was >50 megaohms. The criteria for series resistance and capacitance were ≤20 megaohms and ≥1 picofarad, respectively. A criterion for voltage-clamp stability was defined by a standard error of <10% for baseline current measured at the holding potential for all test pulses. Whole-cell currents were not leak-subtracted. The contribution of background currents was determined by recording before and after addition of the M-current blocker XE-991 (25 μM, Abcam, Cambridge, MA; or TOCRIS, Minneapolis, MN). The currents recorded in XE-991 were digitally subtracted from recordings done in the absence of XE-991. To generate maximum XE-991 block, 8-12 30 s depolarizing voltage pulses to +10 mV were applied at 0.025 Hz before recording. None of the KCNQ2 variants tested in this study exhibited insensitivity to XE-991 under these conditions.

Currents were elicited from a holding potential of -80 mV using 1000 ms depolarizing pulses (from -80 mV to +40 mV in +10mV steps, 0.05 Hz) followed by a 250 ms step to 0 mV to analyze tail currents. Maximum current amplitude was measured 999 ms after the start of each depolarizing voltage pulse whereas tail currents were measured 5 ms after changing the membrane potential to 0 mV. Voltage protocols were repeated in the presence of retigabine (10 µM; Sigma-Aldrich).

### Whole cell patch clamp recording

Manual patch recordings were obtained using an Axon 200B amplifier and pClamp 9 software (Molecular Devices, Sunnyvale, CA) as previously described.^30^ Electroporated cells were thawed and plated onto glass cover slips, then maintained initially at 37°C and subsequently 28°C as described for automated patch clamp experiments. Recordings were obtained using the same internal solution used for automated patch clamp and with internal solutions used previously for manual recordings.^30^ There were no differences in channel behavior between the two internal solutions.

### Data analysis

Data were analyzed and plotted using a combination of DataController384 version 1.8 (Nanion Technologies), Excel (Microsoft Office 2013, Microsoft), SigmaPlot 2000 (Systat Software, Inc., San Jose, CA USA) and Prism 8 (GraphPad Software, San Diego, CA). Whole-cell currents were normalized for membrane capacitance and results expressed as mean ± SEM. The voltage-dependence of channel activation was determined only for cells with mean current density values greater than the background current amplitude. Tail currents were normalized to maximal (peak) tail current amplitude and expressed as a function of the preceding voltages. Data were fit to a Boltzmann function I(V) = Imax/(1 + exp((1/*k*)-(V-V½))), where Imax is the maximum tail current amplitude at test potential V, V½ is the half-maximal activation potential, and *k* is a slope factor. The time-constant of activation (τ) was determined by fitting currents elicited by voltage steps between -30 mV and +40 mV (50-1000 ms after start of the voltage step) to a single exponential function: *A*(*t*) = *A*_0_ × exp(-*t*/τ) + C, where *A*_0_ is the signal amplitude, *t* is time in ms, τ is the time constant of activation, and C is the signal offset.

Typical experiments compared five KCNQ2 variants to the WT channels assayed on the same plate with up to 64 replicate recordings. Properties of each KCNQ2 variant are presented relative to the WT channel assayed in parallel as percent current density measured at +40 mV, difference (Δ) in voltage-dependence of activation V½, and the ratio of activation time-constants (variant / WT). The number of cells used for each experimental condition is given in the figure legends or in Supplemental Tables.

### Statistical analysis

Statistical analysis was performed with one way ANOVA (2 or more variants) or *t*-test, and P ≤ 0.02 was considered significant.

### Data availability

The authors confirm that the data supporting the findings of this study are available within the article and its Supplementary material. Any additional raw data are available on request from the corresponding author.

## RESULTS

### Variant curation and prioritization for functional screening

We assembled a de-identified panel of disease-implicated *KCNQ2* variants from databases of individuals with early-onset epilepsy and or neurodevelopmental impairment participating in research, individuals undergoing clinical genetic testing, and variants described in prior publications. Variants selected for high-throughput functional analysis included “benchmark variants” (recurrent variants with well-established phenotype associations and functional consequences previously determined by voltage-clamp recordings in heterologous cells), variants with established pathogenicity (recurrent *de novo* variants in affected individuals with characteristic *KCNQ2*-associated phenotypes) but without previous functional analysis, and rare variants with less certain pathogenicity. Rare variants present in gnomAD with minor allele frequencies higher than expected for pathogenic variants were included as a presumed non-pathogenic control group. The complete variant panel with associated phenotypes, channel domain, population allele frequency, database and literature references (where applicable) is provided in **Table S1**, and location with the KCNQ2 channel protein for each variant is illustrated in **Fig. S1**.

### Validation of automated patch clamp for assessing KCNQ2 variants

To enhance reproducibility in studies of KCNQ2/KCNQ3 (Q2/Q3) heteromeric channels, we generated a Chinese hamster ovary cell line that stably expresses wild type (WT) human KCNQ3 (CHO-Q3 cells), and then used electroporation with combinations of WT and variant KCNQ2 plasmids to express KCNQ2/KCNQ3 (Q2/Q3) heterotetramers. Automated patch clamp recording demonstrated that maximal (peak) current density measured at +40 mV from CHO-Q3 cells electroporated with WT KCNQ2 was ∼20-fold larger than that measured from non-transfected cells, or from parental CHO cells lacking KCNQ3 but electroporated with KCNQ2 (**Fig. 1A-C**). The lack of current obtained from cells expressing KCNQ2 or KCNQ3 alone suggested that currents from KCNQ2 electroporated CHO-Q3 cells were likely Q2/Q3 heterotetramers. Indeed, the sensitivity of the measured currents to tetraethylammonium (TEA) ion (IC_50_, 10.7 ± 4.2 mM; **Fig. 1D**) is consistent with previous reports of TEA sensitivity for Q2/Q3 channels expressed in CHO cells, and unlike the sensitivity or either homotetrameric KCNQ2 (0.2 mM) or KCNQ3 (>100 mM) channels.^31,32^ Currents recorded from the same cells by conventional manual patch clamp exhibited similar voltage-dependent gating properties (**Fig. S2**). Differences in current density between manual and automated methods appeared related to differences in sampling (e.g., use of GFP expression to select higher expressing cells for manual patch clamp recording). Finally, we tested the effects of retigabine on Q2/Q3 channel activity (**Fig. 2**). Retigabine (10 μM) induced a 1.4-fold increase in peak current density measured at +40 mV accompanied by a strong hyperpolarizing shift (V½ shifted -29.2 ± 0.3 mV) in the voltage-dependence of activation and faster activation kinetics in the -30 to -10 mV range, in agreement with previous reports.^32-35^

**Figure 1.**
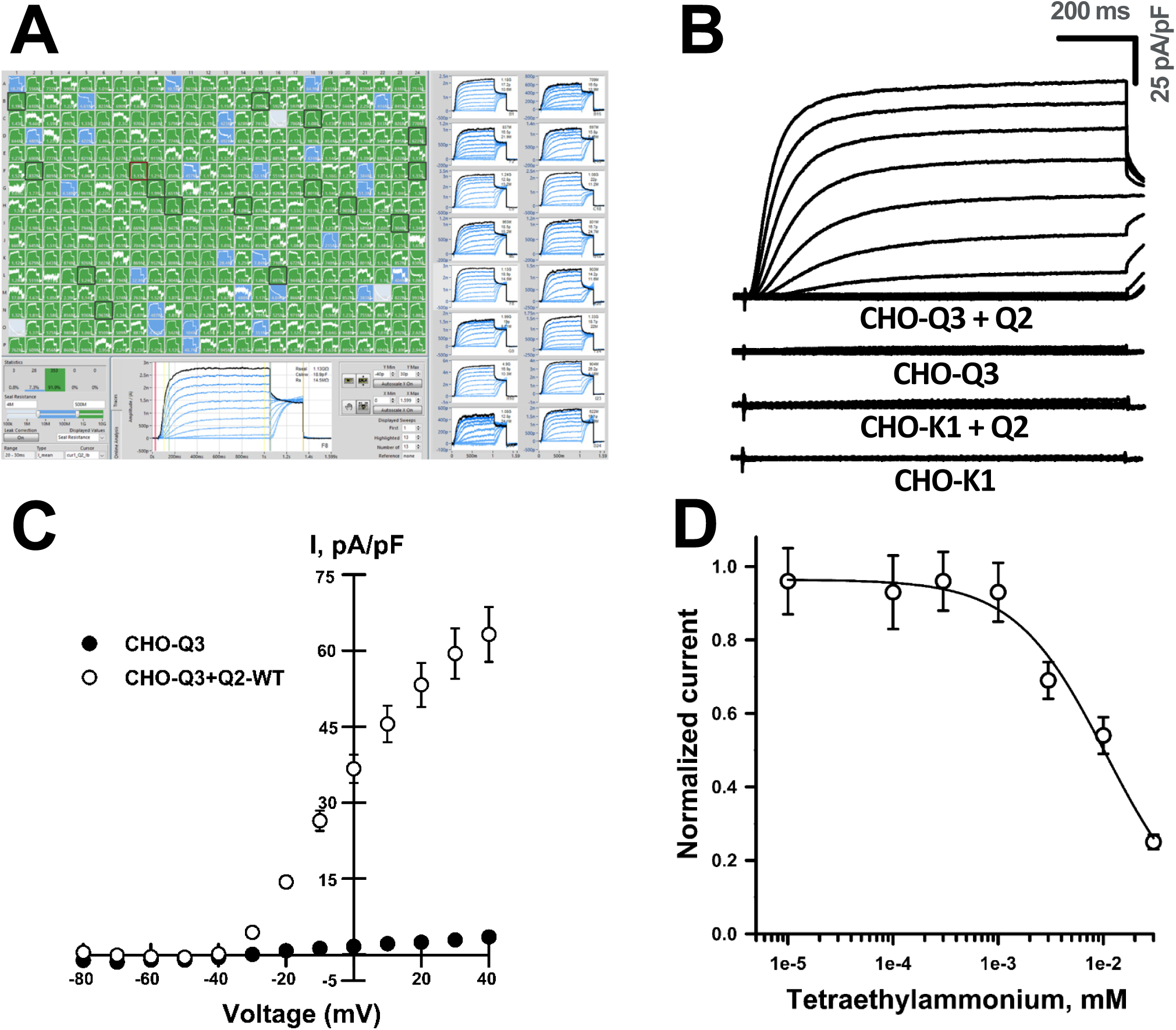
Functional analysis of KCNQ2/KCNQ3 channels by automated patch clamp. **A**) Screen display from automated patch clamp experiment illustrating whole-cell current recordings from CHO-Q3 cells transiently expressing KCNQ2 variants (6 variants, 4 columns per variant). **B**) Averaged XE-991-sensitive whole-cell currents recorded from non-transfected CHO-K1 cells, CHO-K1 cells electroporated with WT KCNQ2, non-transfected CHO-Q3 cells, and CHO-Q3 cells electroporated with WT KCNQ2. **C**) Average current-voltage relationships measured from non-transfected (open circles, n = 94) or KCNQ2-WT transfected (filled circles, n = 124) CHO-Q3 cells. Current recorded from each cell was normalized to cell capacitance as a surrogate for cell size to calculate current density (I, pA/pF). **D**) Concentration-response relationship for TEA block of whole-cell currents recorded from CHO-Q3 electroporated with KCNQ2-WT (IC_50_ = 10.7 ± 4.2 mM, n = 51-82 for each concentration). Horizontal and vertical scale bars represent 200 ms and 20 pA/pF, respectively.

**Figure 2.**
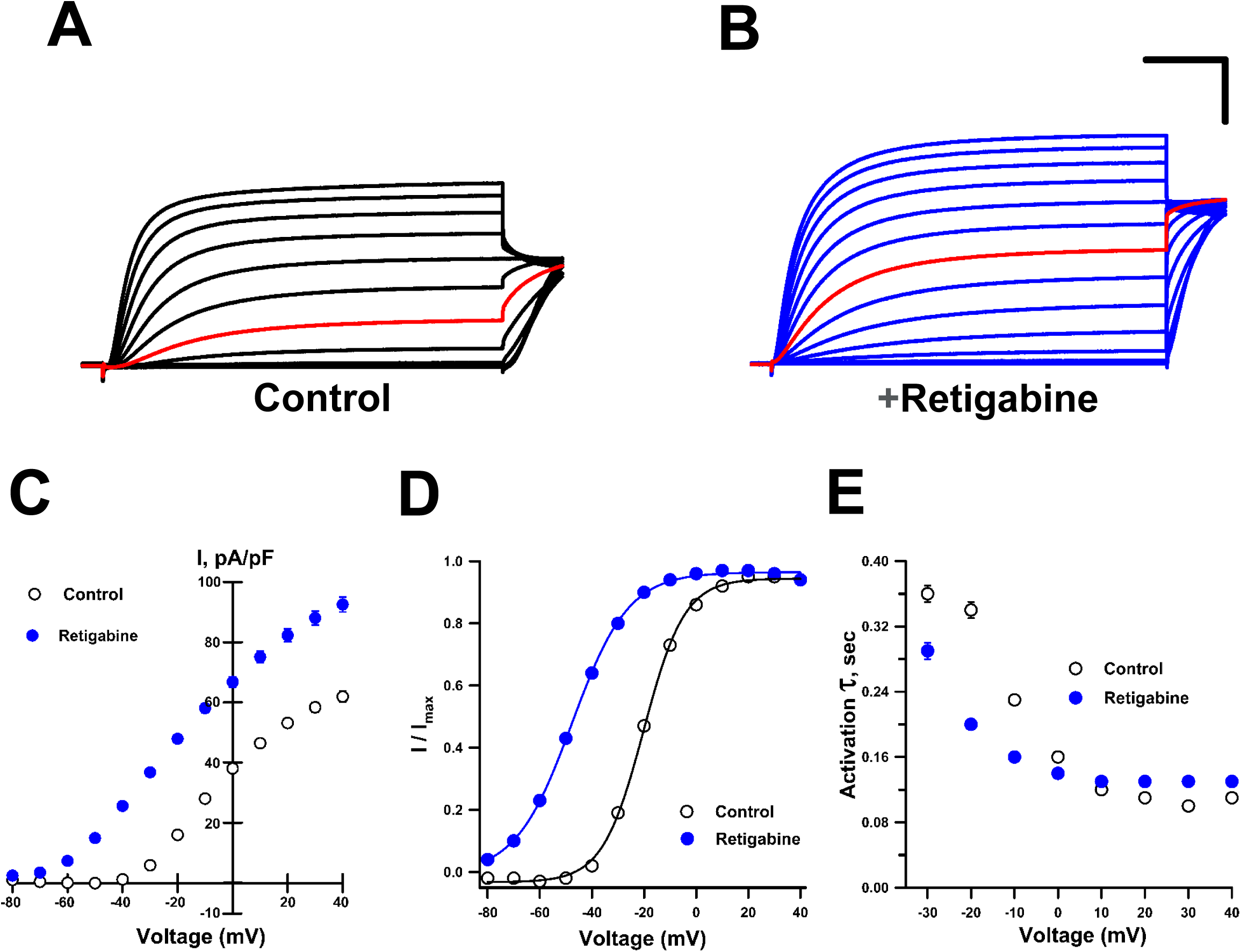
Effects of retigabine on KCNQ2/KCNQ3 channel activity. Averaged XE-991-sensitive whole-cell currents normalized by membrane capacitance recorded from CHO-Q3 cells electroporated with KCNQ2-WT exposed to control (**A**) or 10 μM retigabine (**B**) solutions. Red lines indicate currents recorded at -20 mV. **C**) Average current-voltage relationships measured from CHO-Q3 cells electroporated with KCNQ2-WT exposed to control (open circles, n = 1086) or retigabine (filled blue circles, n = 1141) solutions. **D**) Voltage-dependence of activation measured in control (open circles, black lines) or retigabine (filled blue circles, blue lines) solutions (control: V½ = -18.9 ± 0.2, *k* = 7.6 ± 0.1, n = 833; retigabine: V½ = - 47.7 ± 0.3, *k* = 9.9 ± 0.1, n = 885). **E**. Activation time constants measured in control (open circles, n =525-970) or retigabine (filled blue circles, n = 1036-1079) solutions. Horizontal and vertical scale bars represent 200 ms and 20 pA/pF, respectively.

To demonstrate that automated patch clamp recording is a valid approach for evaluating the functional consequences of epilepsy-associated *KCNQ2* variants, we compared data we obtained for 18 variants to results from prior voltage-clamp recording in either mammalian cells or *Xenopus* oocytes.^11,14,18,20-22,25,36-41^ To enable comparison of channel properties resolved by different experimental approaches, we quantified differences in peak current density and the midpoint of activation voltage dependence (V½) relative to WT channels assayed in parallel. Low current densities of KCNQ2 homomeric channels (i.e., absence of KCNQ3) in our system precluded study of this channel type. Instead, we either electroporated each variant alone in the CHO-Q3 cells, mimicking homozygosity, or electroporated a 1:1 ratio of WT and variant KCNQ2, mimicking heterozygosity. We then compared the results obtained for each variant with prior work in the homozygous (**Fig. 3**) and heterozygous (**Fig. S5**) states (complete data set presented in **Table S3**). Averaged whole-cell currents normalized to peak current amplitude measured for WT channels were recorded from literature variants in the homozygous (**Fig.S3**) and heterozygous (**Fig.S4**) configurations. Results were concordant between automated and manual patch clamp recording for the majority of the 18 literature variants despite the diversity of laboratories and expression systems (**Fig. 3**). This included variants with loss-of-function and gain-of-function effects. There were divergent results for only 2 BFNE variants (peak current density for R333Q in the homozygous state [**Fig. 3B**] and L243F in the heterozygous state [**Fig. S5A**]), and small quantitative differences for a small number of other variants. We concluded that automated patch clamp recording was a robust and valid strategy to assess the functional properties of KCNQ2 variants.

**Figure 3.**
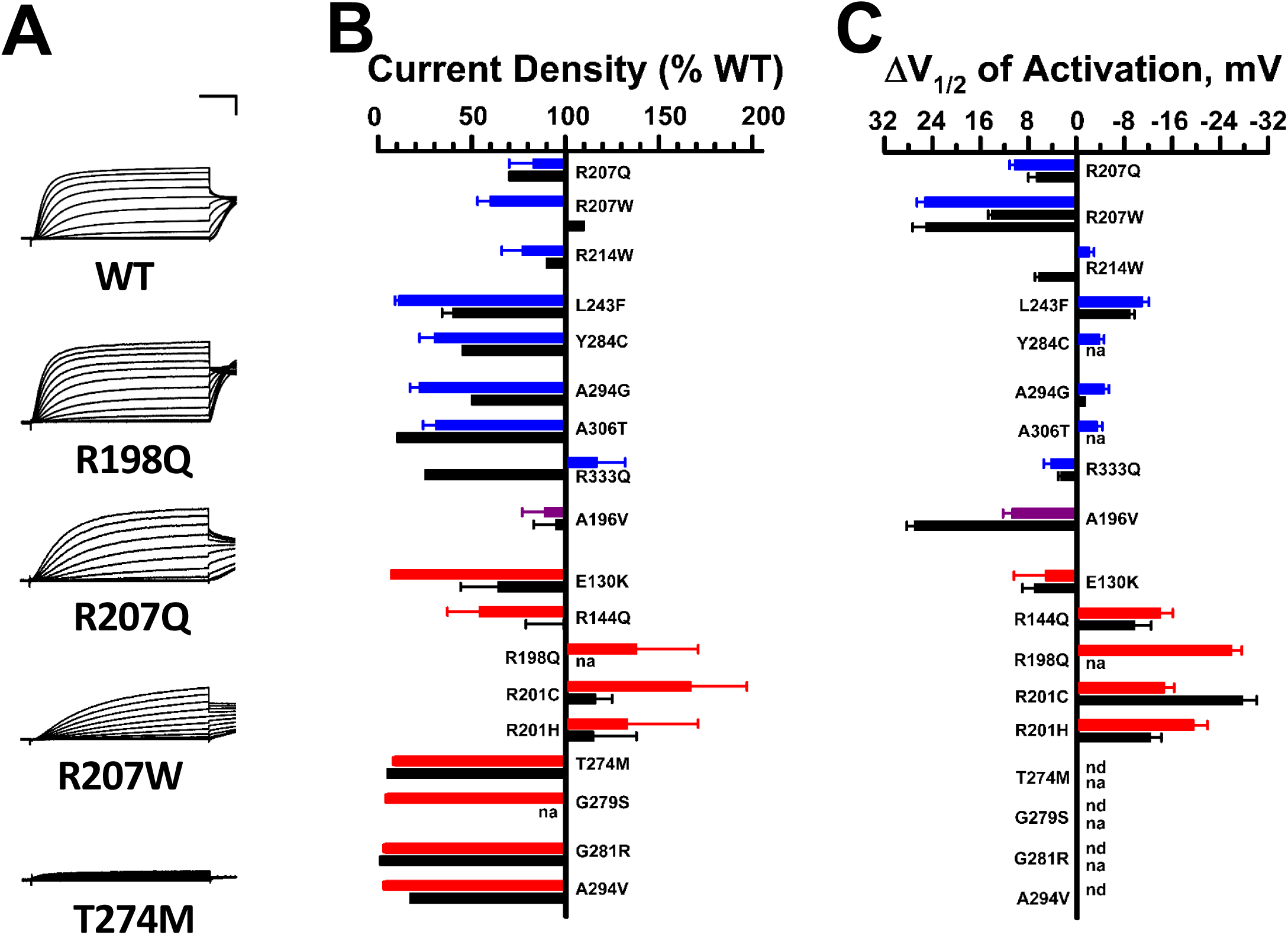
Functional properties of homozygous KCNQ2 variants determined by automated patch recording are comparable to those from previous voltage-clamp studies. **A)** Examples of averaged XE-991-sensitive whole-cell currents recorded by automated patch clamp from CHO-Q3 cells electroporated with select KCNQ2 variants. Current values were normalized to WT channel peak current that was measured in parallel. **B)** Average peak whole-cell currents recorded at +40 mV from cells expressing KCNQ3 plus KCNQ2 variants displayed as percent of WT channel that was measured in parallel. **C)** Difference in activation V½ determined for KCNQ3 plus KCNQ2 variant channels relative to WT channel (horizontal scaling was designed to show loss-of-function in the leftward direction from zero). Horizontal and vertical scale bars represent 200 ms and 25%, respectively. Black bars indicate literature voltage-clamp data, while automated patch clamp results are shown as blue bars for BFNE, red bars for DEE, and purple bars for BFNE/DEE pathogenic variants. Values that were not available in the literature are indicated by na, and values that could not be determined are indicated by nd.

### Functional analysis of rare population variants

To assess KCNQ2 functional variation in the general population, we studied 24 population variants obtained from the gnomAD database^42^ (**Table S2**). Most of these variants are localized to cytosolic domains (17 C-terminus, 1 N-terminus) whereas 3 variant residues lie within transmembrane segments or an intersegment linker region (S1-S2). Averaged whole-cell currents for the population variants expressed in the homozygous state are shown in **Fig. S6**, and differences in peak current density or voltage dependence of activation V½ compared with WT channels are summarized in **Fig. 4** (complete data set presented in **Table S4**). No variant showed large differences in gating kinetics, but 5 variants (I238V, P410L, A503V, R604C and T771I) exhibited significantly smaller peak current densities (40-60%, **Fig. 4A**) compared to the WT channel, and 8 variants had significant (≥5 mV) shifts in activation V½ in either the depolarizing (loss of function: F104L and T605S) or hyperpolarizing (gain of function: P410L, A503V, S751L, R760H, N780T and R854C) direction (**Fig. 4B**; **Table S5**). However, when expressed in the heterozygous state, none of the population variants exhibited significantly lower current density or significant shifts in activation V½ (**Fig. 4C,D; Fig. S7**). However, two rare population variants (F701del, S751L) had significantly greater peak current densities when studied in the heterozygous state. These data provide evidence that in the general population, loss-of-function is unusual with heterozygous KCNQ2 variants.

**Figure 4.**
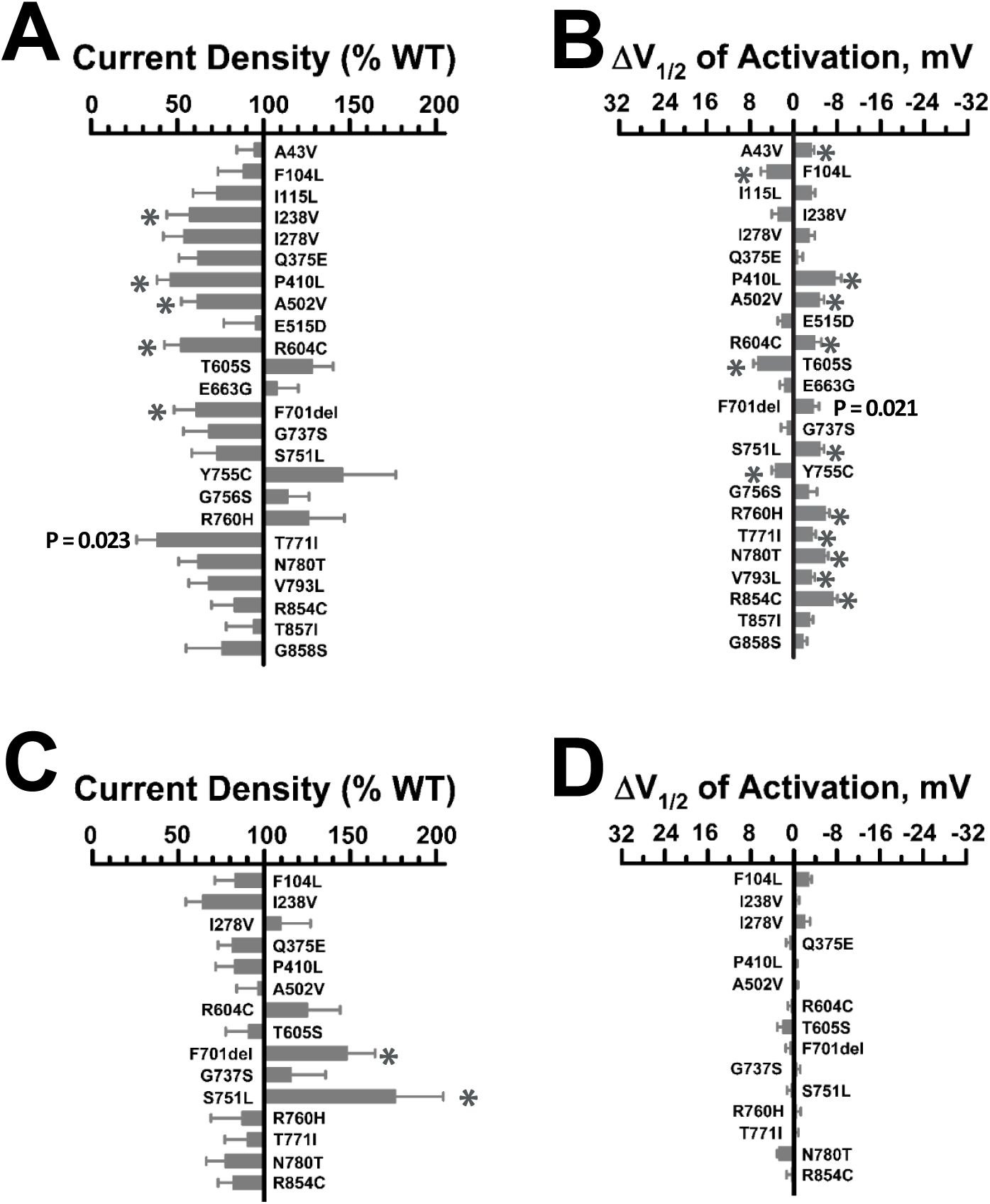
Functional properties of KCNQ2 population variants. **A**) Average whole-cell currents recorded at +40 mV expressed as percent of the WT channel (n = 22-71). Variants were expressed in the homozygous state in CHO-Q3 cells. **B**) Differences in activation V½ (n = 15-60) relative to the WT channel determined for KCNQ2 population variants. Full datasets are provided in **Table S4. C**) Average whole-cell currents recorded at +40 mV (n = 27-71) for select population variants expressed in the heterozygous state in CHO-Q3 cells. Only variants exhibiting significantly different properties from WT in the homozygous state were examined. **D**) Differences in activation V½ (n = 9-59) relative to WT channel determined for select population variants expressed in the heterozygous state. Full datasets are provided in **Table S5**. Statistical significance (*p* ≤ 0.02) is indicated by single asterisks (^*^).

### Functional analysis of epilepsy-associated KCNQ2 variants

We determined the functional properties of 39 additional epilepsy-associated KCNQ2 variants that had not been investigated at the time of our experiments (8 BFNE, 30 DEE, 1 unknown phenotype; **Fig. S1, Table S2**). Averaged whole-cell currents recorded from Q3-CHO cells expressing each variant in the homozygous state are presented in **Fig. S8**, and differences in peak current density or voltage dependence of activation V½ relative to the WT channel are summarized in **Fig. 5** (complete data set presented in **Table S4**). The majority of the variants exhibited severe loss-of-function (<25% peak current density compared with WT channels) with profound loss-of-function (≤10% of WT) observed for 18 variants. The remaining variants exhibited less severe degrees of dysfunction (peak current density 25-75% of WT) or peak current density that was not significantly different from WT. None of the 39 variants exhibited current density larger than WT channels. For some variants with tail currents large enough to determine the voltage-dependence of activation, we observed significant differences from WT channels with shifts in activation V½ values in either the depolarizing (loss-of-function) or hyperpolarizing (gain-of-function) direction (**Fig. 5B**). A depolarized activation V½ was observed for variants Q204H, H228Q, S113F, L203P, R210H, whereas variants V543M, M578V, R144W, A193D, Q586P exhibited hyperpolarized shifts in V½. In addition, the time course of activation was not affected by most variants with the exception of S113F and A193D, which had 3-fold and 1.5-fold slower activation, respectively (see activation time constants in **Table S4**; **Fig. S8**). Only the R333W variant was not significantly different from WT channels in any functional property using a stringent threshold for significance. Importantly, there were no systematic differences in the functional properties of the tested missense variants associated with BFNE or DEE.

**Figure 5.**
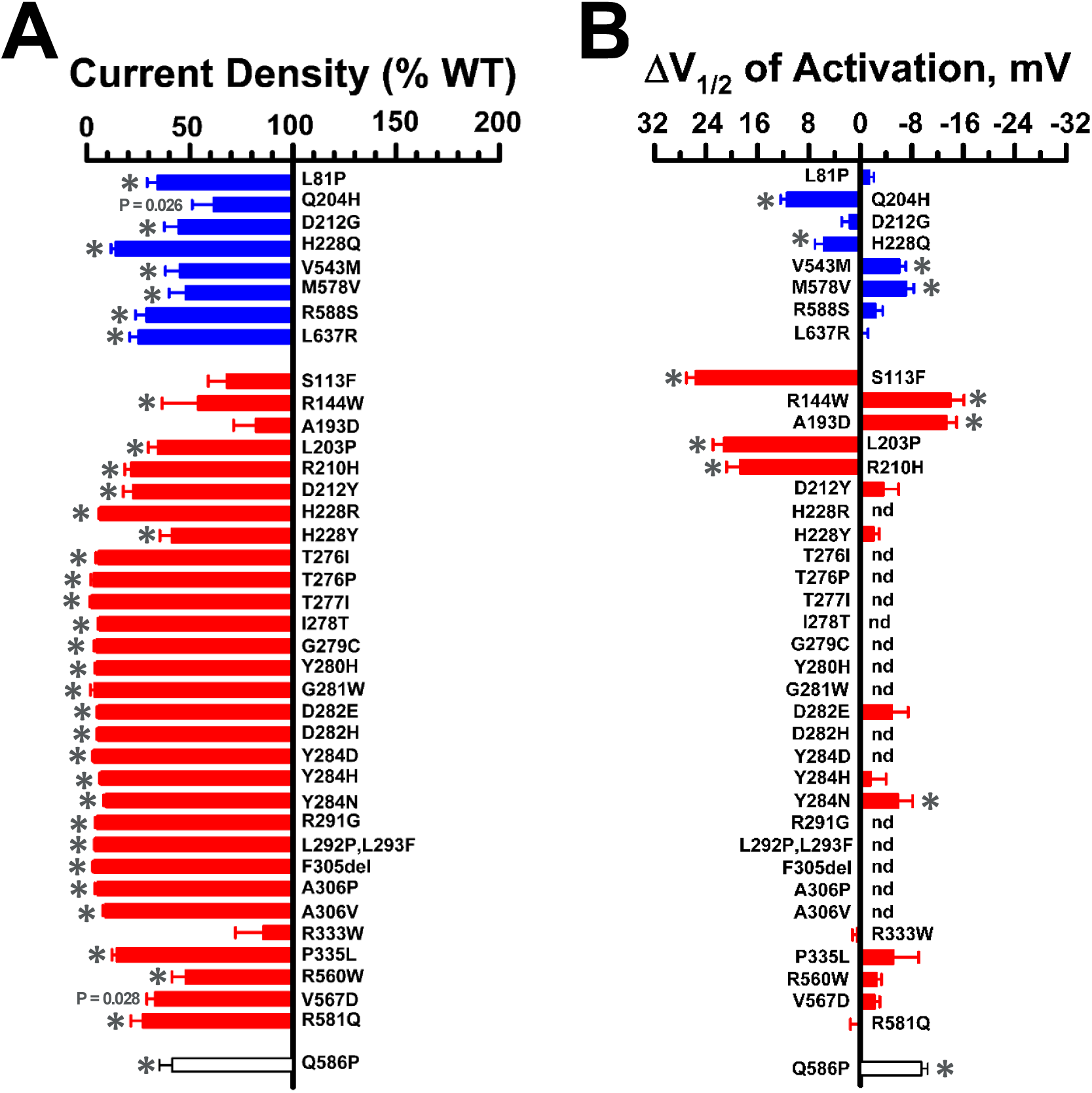
Functional properties of epilepsy-associated KCNQ2 variants studied in the homozygous state. **A**) Average whole-cell currents recorded at +40 mV from CHO-Q3 cells electroporated with epilepsy-associated KCNQ2 variants displayed as percent of WT channel (n = 16-75). **B**) Differences in activation V½ determined for disease-associated variant channels relative to the WT channel (n = 7-41). Blue bars indicate variants associated with BFNE, red bars are variants associated with DEE, the purple bar is a variant associated with both phenotypes, and the uncolored bar is a variant with an unclear phenotype. Values that could not be determined are indicated by nd. Statistical significance (*p* ≤ 0.02) is indicated by single asterisks (^*^). Full datasets are provided in **Table S4**.

Because pathogenic KCNQ2 variants associated with epilepsy are heterozygous, we assessed the functional properties of each variant expressed in this context by co-transfecting Q3-CHO cells with equal amounts of WT and variant KCNQ2 plasmids. Averaged whole cell currents recorded in these experiments are presented in **Fig. S9** and the differences in peak current density or activation V½ relative to the WT channel are summarized in **Fig. 6** (complete data set presented in **Table S5**). Dominant-negative behavior, which we defined as peak current density less than 50% of WT heterotetrameric channels, was observed for several of the variants we studied and was more prevalence among DEE then BFNE variants. Some variants exhibited significantly lower current density than WT channels but to a lesser degree than dominant-negative variants (V543M, H228Y, G281W, Y284H) and this is more consistent with haploinsufficiency. Most of the remaining variants showed no difference in peak current density compared with WT channels and one variant (Q586P) exhibited larger current density. As observed in the homozygous state, many variants exhibited significant shifts in activation V½ relative to the WT channel, and some of the variants that lacked a significantly different peak current density exhibited significant shifts in activation V½ relative to the WT channel (R144W, P335L, V567D). Three variants (Q204H, H228Q, R333W) exhibited no significant differences in peak current density or activation V½ in the heterozygous state relative to the WT channel.

**Figure 6.**
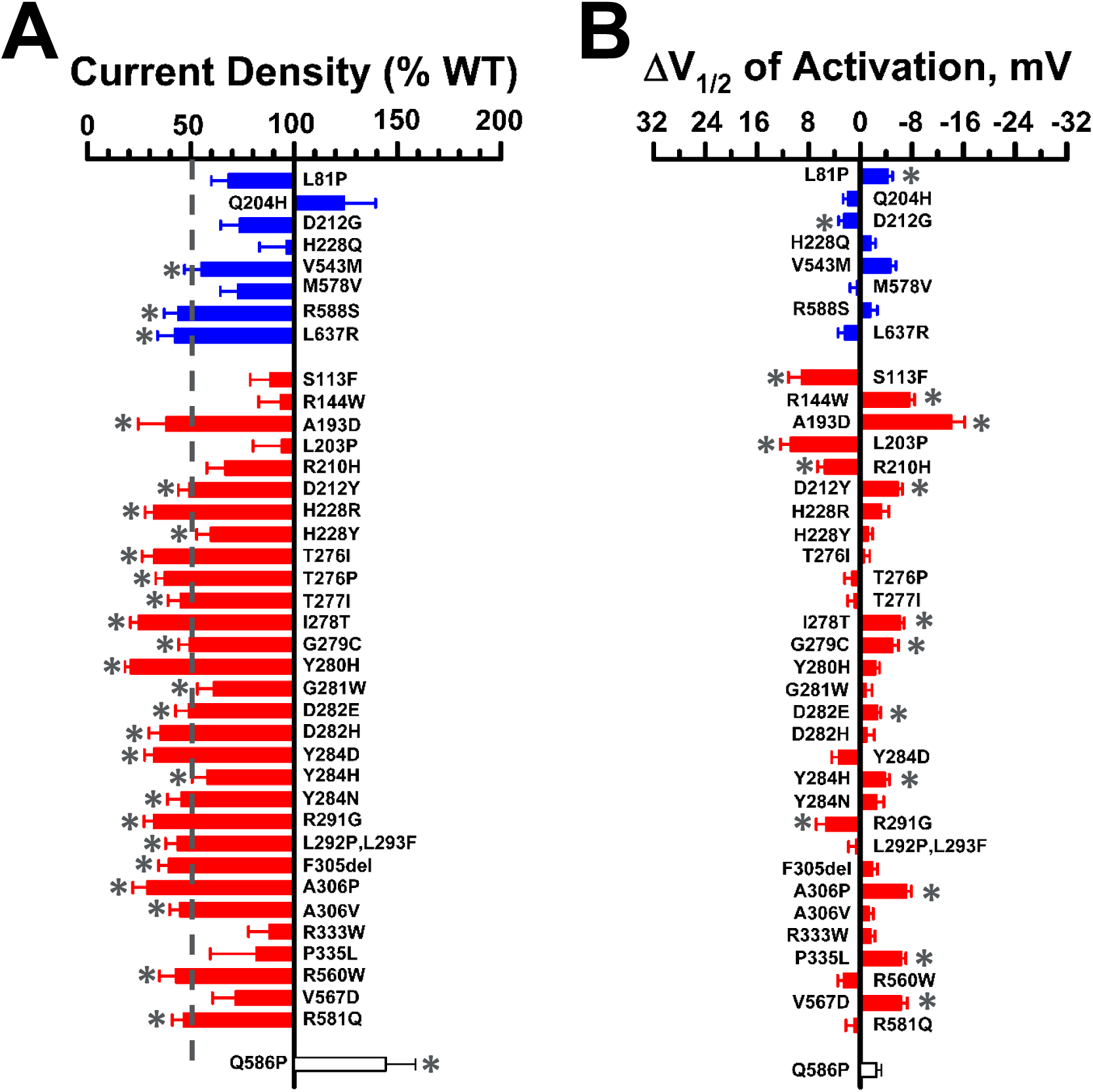
Functional properties of epilepsy-associated KCNQ2 variants studied in the heterozygous state. Average whole-cell currents recorded at +40 mV from CHO-Q3 cells co-electroporated with WT and epilepsy-associated variant cDNAs and displayed as percent of WT channel (n = 19-84). Differences in activation V½ determined for epilepsy-associated KCNQ2 variants relative to the WT channel (n = 13-69). Blue bars indicate variants associated with BFNE, red bars are variants associated with DEE, the purple bar is a variant associated with both phenotypes, and the uncolored bar is a variant with an unclear phenotype. Statistical significance (*p* ≤ 0.02) is indicated by single asterisks (^*^). Full datasets are provided in **Table S5**.

### Effects of retigabine on KCNQ2 variants

To assess the pharmacological consequences of KCNQ2 variants, we tested the acute effects of the positive allosteric modulator retigabine (10 μM) on channel function. When variants were expressed in the homozygous state, we observed a wide range of retigabine effects on maximal current density and activation V½ (**Fig. S10**; complete data set in **Table S6**). Some variants exhibited little to no response, whereas others showed large effects equal to or greater than WT channels. Some variants located in the S4 segment and S4-5 linker (R198Q, H228Q, H228Y) exhibited a partial response characterized by a substantial hyperpolarizing shift in activation V½ but without a WT-like increase in current density. For S4 variants R207W and R207Q, which cause a depolarized shift of activation V½ and a marked slowing of channel opening rates across voltages (**Fig. 3A-C**), retigabine countered both effects resulting in a greater relative increase in current density than observed for WT channels. Variants largely unresponsive to retigabine involved residues between 280 and 306, which encompasses portions of the pore domain from the ion selectivity filter to the nearby extracellular half of the S6 segment. These data provide a map of residues with strong effects on ion conduction that cannot be overcome by retigabine when expressed in the homozygous context.

Because retigabine is a potential therapeutic agent for KCNQ2-associated epilepsy, we investigated its effect on epilepsy-associated variants expressed in the heterozygous state (**Table S7**). We compared current density measured across a range of voltages for each heterozygous variant in the presence of retigabine with that of the WT channel recorded in the absence of drug (**Fig. S11**). Retigabine restored current density to at least the level of untreated WT channels for all epilepsy-associated variants at voltages more negative than the activation V½ of WT channels. However, as predicted by results from studying homozygous channels, the effect of retigabine on current density at strongly depolarized voltages varied considerably among variants. Variants affecting residues within the ion selectivity filter (TIGYG) that exerted strong dominant-negative effects (e.g., Y280H, A306T; **Fig. 7A**) exhibited the smallest responses to retigabine throughout the voltage range (**Fig. S11; Table S7**). Several well-established gain-of-function variants within the voltage-sensor domain (R144Q, R198Q, R201H) and others elsewhere in the channel were hyper-responsive to retigabine (examples shown in **Fig. 7A; Fig. S11; Table S7**). A summary of effects of retigabine on current density measured at -20 mV for all epilepsy-associated variants is presented as a heat map in **Fig. 7B** illustrating that *in vitro* responses to retigabine exhibit considerable heterogeneity among variants.

**Figure 7.**
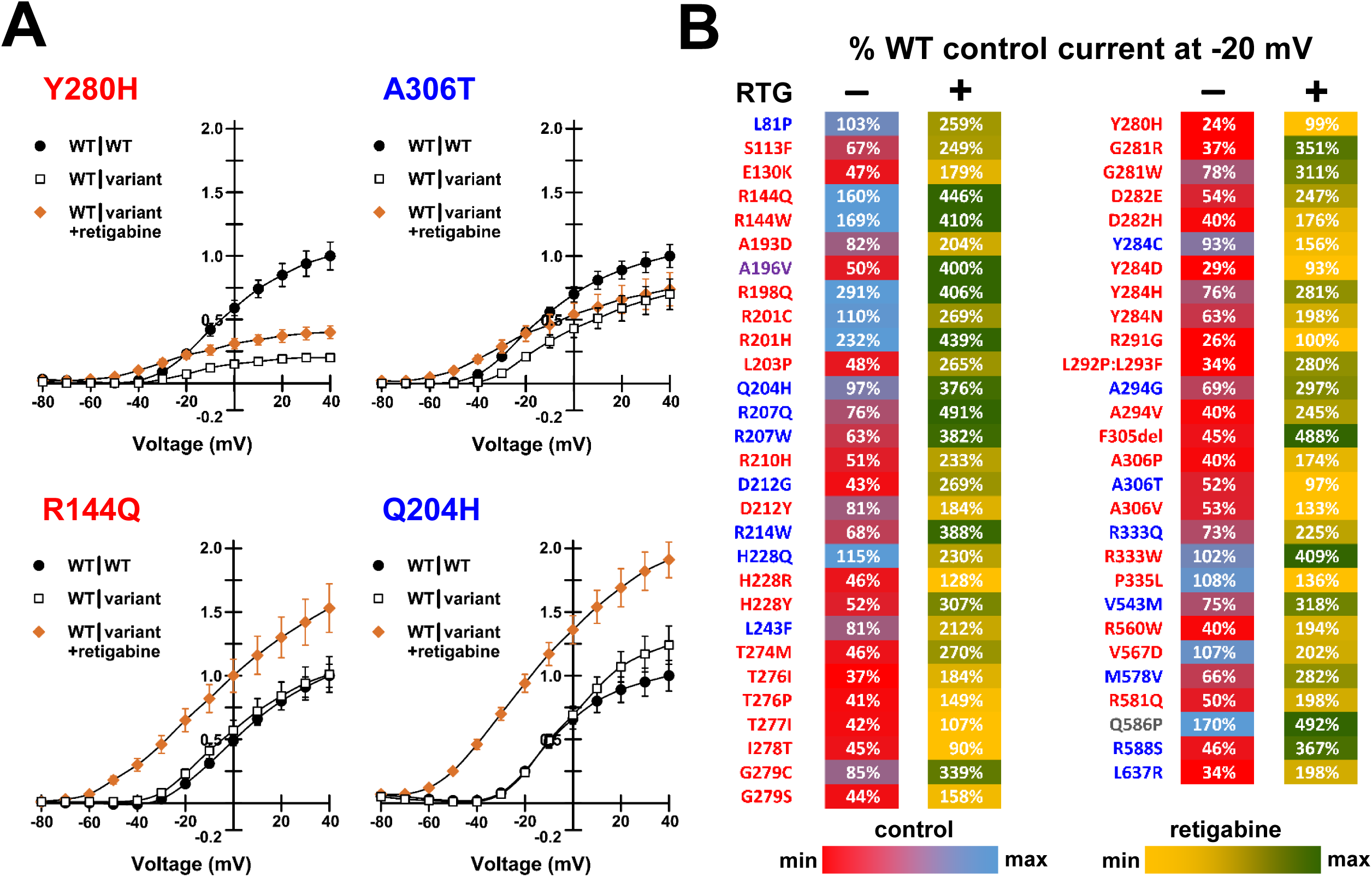
Responses of epilepsy-associated KCNQ2 variants to retigabine. **A**) Representative current-voltage relationships comparing WT (WT|WT, filled circles) and heterozygous variant (WT|variant, open squares) channel function in the absence of retigabine with heterozygous variant function measured in the presence of 10 µM retigabine (WT|variant +retigabine, orange filled diamonds). Current amplitude was first normalized to cell capacitance, then normalized to the WT current density measured at +40 mV. Variant labels are color coded by phenotype (blue, BFNE; red, DEE; purple, BFNE/DEE; gray, unclear phenotype). **B**) Heat map summarizing retigabine response data. Control values are current density measured at -20 mV for heterozygous variants in the absence of retigabine expressed as a percentage of untreated WT channel current density (**Table S5**). Retigabine values are current density measured at -20 mV for heterozygous variants in the presence of 10 µM retigabine and expressed as a percentage of untreated WT channel current density (**Table S7**). Each value is colored based on the scale shown.

## DISCUSSION

Advances in understanding monogenic forms of epilepsy has motivated an aspirational approach in which treatments are designed to target specific genes or specific genetic variants.^43^ Before the potential of this precision medicine paradigm can be realized, a more comprehensive understanding of the pathophysiological consequences of genetic variants is needed. This requires expanding the throughput and capacity for evaluating the functional and pharmacological properties of hundreds, or even thousands, of variants using well-validated approaches, then implementing these experimental platforms for assessing genotype-specific drug responses. Additionally, incorporating data from functional studies into variant classification schemes will improve the diagnostic accuracy of genetic testing.

In this study, we undertook the largest single effort to determine the functional and pharmacological properties of epilepsy-associated *KCNQ2* variants in both the homozygous and heterozygous state. The unprecedented scale of this work was made possible by using automated patch clamp recording rather than traditional voltage clamp methods, which have limited throughput. Automated patch clamp has specific advantages over manual methods including higher throughput, unbiased cell selection, and efficient coupling to pharmacological studies. The higher throughput enables larger numbers of replicates, which strengthens statistical power and experimental rigor.

Demonstrating deleterious consequences of genetic variants assessed by well-established *in vitro* or *in vivo* methods is considered strong evidence supporting pathogenicity in the framework for classifying variant developed by the American College of Medical Genetics and Genomics (ACMG).^44^ Inclusion of multiple control variants is critically important to ensure reliability for a functional assay.^45^ To validate use of automated patch clamp for KCNQ2, we performed initial experiments on 18 epilepsy-associated variants that were studied previously and 24 population variants that were expected to exhibit non-pathogenic properties. Therefore, our method satisfies these published criteria for a well-established functional assay.

After validating our approach, we investigated 39 epilepsy-associated variants with unknown functional consequences. These included variants with established clinical pathogenicity based on clear segregation in families or recurrence as de novo variants in unrelated individuals, as well as variants with uncertain pathogenicity. We observed that the majority of these KCNQ2 epilepsy-associated variants exhibit properties consistent with loss-of-function that can be predicted to cause impaired neuronal M-current. A small number of the epilepsy-associated variants studied here exhibited functional properties that were not dramatically different from WT channels. There are multiple potential explanations for this finding. As our sample included variants of uncertain significance on clinical grounds, some may not be disease causing and their discovery in persons with epilepsy may be incidental. Although demonstrating normal function in heterologous cells is an important observation, some variants may only have abnormal behavior in neurons. Lastly, some variants have been demonstrated to exhibit their strongest functional effects when expressed in the absence of KCNQ3 (e.g., as KCNQ2 homotetramers).^41^ Unfortunately, we were unable to measure current in cells expressing KCNQ2 only.

Our study uniquely investigated the effects of retigabine on several KCNQ2 variants, and these data allowed us to recognize the variable nature of this drug response among variants. Retigabine was initially approved for adjunctive treatment of partial epilepsy in adults,^32^ but was withdrawn from the market because of limited use in part because of an adverse blue discoloration of skin.^46^ However, the drug may have potential value as targeted therapy for KCNQ2-associated epilepsy.^11^ Retigabine acts to accentuate M-current through altering the voltage-dependence of activation of KCNQ2/KCNQ3 channels. The drug binds to a hydrophobic pocket comprised of residues from both channel subunits including a tryptophan residue at position 236 in KCNQ2 and neighboring residues (e.g., Leu-299, Ser-303, Phe-305) in the three-dimensional structure.^47,48^ None of the variants we studied affect residues known to be involved directly in retigabine binding.

We initially examined the effects of retigabine on KCNQ2 variants expressed in the homozygous state and observed a wide range of responses. These observations may contribute to advancing understanding structure-pharmacology relationships for KCNQ2/KCNQ3 channels. We also conducted experiments to determine the response of epilepsy-associated variants expressed in the heterozygous state to ascertain how retigabine affects channel activity in a clinically relevant context. By comparing current density of retigabine modulated variant channels with unmodulated WT channels at a membrane potential relevant to the physiological contribution of M-current on neuronal excitability, we observed that function of all epilepsy-associated variants we tested could be restored to at least the level of WT channels. In some cases, current density was boosted to levels exceeding WT channels particularly for variants exhibiting gain-of-function effects in the absence of drug. However, there were large differences in the effects of retigabine on heterozygous channel current density at more depolarized potentials. The correlation between these *in vitro* data and the clinical response to retigabine remains to be determined.

In summary, we performed a large scale *in vitro* functional and pharmacological evaluation of KCNQ2 variants in the homozygous and heterozygous states using automated patch clamp recording. Our findings contribute to understanding the spectrum of KCNQ2 channel dysfunction in monogenic epilepsy and reveal variable retigabine responsiveness among the studied variants. We also observed that dominant-negative missense variants are not exclusive to KCNQ2-associated DEE. The validated functional assay we employed in this study provided data that can be used to help distinguish true pathogenic from non-pathogenic variants, and serve as a platform for evaluating drug responses of other KCNQ2 variants.

## ABBREVIATIONS

BFNE: benign familial neonatal epilepsy
DEE: developmental and epileptic encephalopathy
PV: population variant
Q2: KCNQ2 channel
Q3: KCNQ3 channel
CHO-Q3: cell line stably expressing KCNQ3
V½: voltage at 50% activation
TEA: tetraethylammonium
EGFP: enhanced green fluorescent protein
WT: wild-type

## ACKNOWLEDGEMENTS

The authors thank Jean-Marc DeKeyser for help with molecular biology and for creating the image in Fig. S1. This work was supported by NIH grant NS108874.

## SUPPLEMENTARY MATERIAL

### Supplemental Figures

Fig. S1 KCNQ2 variants analyzed in this study

Fig. S2 Comparison of automated and manual patch clamp recording of KCNQ2/KCNQ3

Fig. S3 Whole-cell currents from literature KCNQ2 variants (homozygous state)

Fig. S4 Whole-cell currents from literature KCNQ2 variants (heterozygous state)

Fig. S5 Manual and automated patch clamp analyses of KCNQ2 variants (heterozygous state)

Fig. S6 Whole-cell currents of KCNQ2 population variants (homozygous state)

Fig. S7 Whole-cell currents of KCNQ2 population variants (heterozygous state)

Fig. S8 Whole-cell currents of KCNQ2 epilepsy variants (homozygous state)

Fig. S9 Whole-cell currents of KCNQ2 epilepsy variants (heterozygous state)

Fig. S10 Retigabine effects on KCNQ2 variants expressed in the homozygous state

Fig. S11 Retigabine effects on KCNQ2 variants expressed in the heterozygous state

### Supplemental Tables

Table S1 KCNQ2 variant information

Table S2 Sequence of mutagenic primers used to generate KCNQ2 variants

Table S3 Data from manual and automated patch clamp recording of KCNQ2 in CHO-Q3 cells

Table S4 Functional properties of homozygous KCNQ2 variants under control conditions

Table S5 Functional properties of heterozygous KCNQ2 variants under control conditions

Table S6 Functional properties of homozygous KCNQ2 variants after exposure to retigabine

Table S7 Functional properties of heterozygous KCNQ2 variants after exposure to retigabine

