## Supplementary Figures and Tables for "High-throughput Evaluation of Epilepsy-associated *KCNQ2* Variants Reveals Functional and Pharmacological Heterogeneity"

### SUPPLEMENTARY MATERIAL

*Supplemental Figures*

Fig. S1 KCNQ2 variants analyzed in this study

Fig. S2 Comparison of automated and manual patch clamp recording of KCNQ2/KCNQ3

Fig. S3 Whole-cell currents from literature KCNQ2 variants (homozygous state)

Fig. S4 Whole-cell currents from literature KCNQ2 variants (heterozygous state)

Fig. S5 Manual and automated patch clamp analyses of KCNQ2 variants (heterozygous state)

Fig. S6 Whole-cell currents of KCNQ2 population variants (homozygous state)

Fig. S7 Whole-cell currents of KCNQ2 population variants (heterozygous state)

Fig. S8 Whole-cell currents of KCNQ2 epilepsy variants (homozygous state)

Fig. S9 Whole-cell currents of KCNQ2 epilepsy variants (heterozygous state)

Fig. S10 Retigabine effects on KCNQ2 variants expressed in the homozygous state

Fig. S11 Retigabine effects on KCNQ2 variants expressed in the heterozygous state

*Supplemental Tables*

Table S1 KCNQ2 variant information

Table S2 Sequence of mutagenic primers used to generate KCNQ2 variants

Table S3 Data from manual and automated patch clamp recording of KCNQ2 in CHO-Q3 cells

Table S4 Functional properties of homozygous KCNQ2 variants under control conditions

Table S5 Functional properties of heterozygous KCNQ2 variants under control conditions

Table S6 Functional properties of homozygous KCNQ2 variants after exposure to retigabine

Table S7 Functional properties of heterozygous KCNQ2 variants after exposure to retigabine

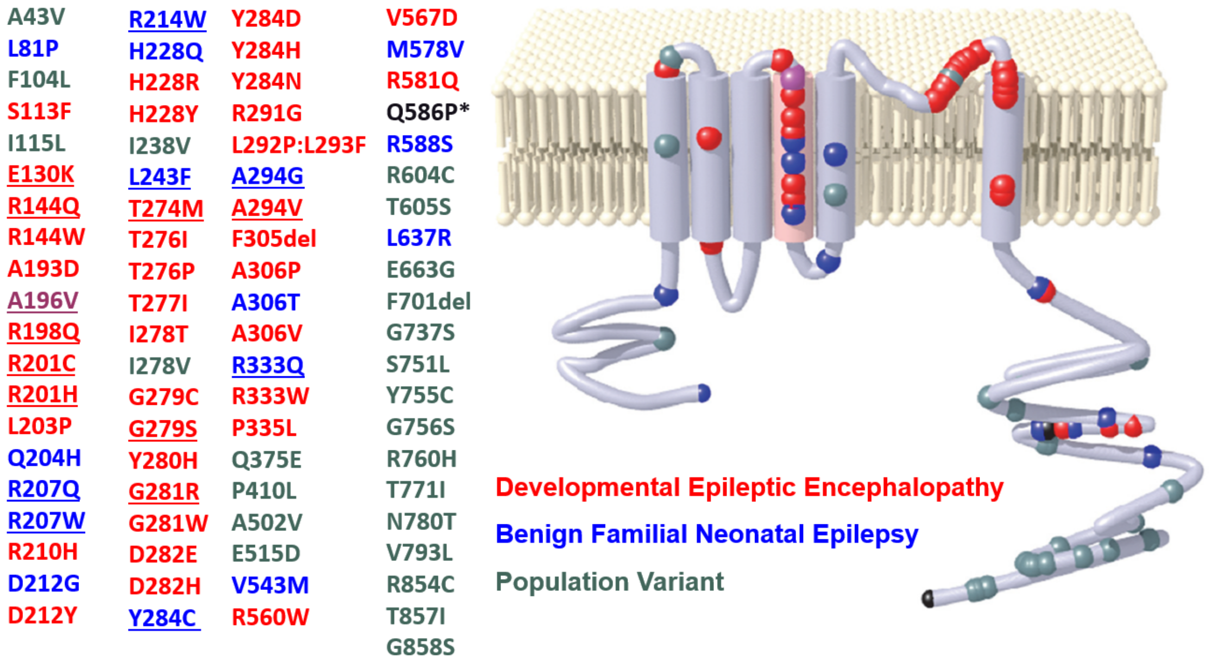


**Figure S1. KCNQ2 variants analyzed in this study.** Location and classification of the 81 KCNQ2 variants analyzed in this study. BFNE-associated variants are shown as blue dots, DEE-associated variants as red dots, the purple dot represents a variant associated with both BFNE and DEE, and population variants are denoted as green dots. Variant Q586P (marked by *) is associated with unknown phenotype category. Literature variants are underlined.

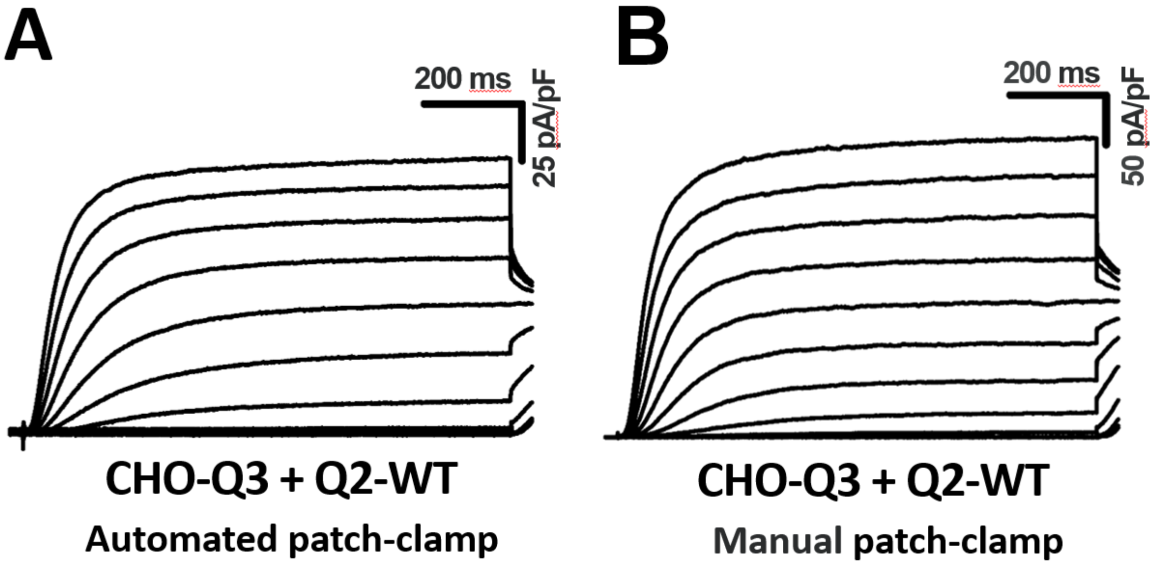


**Figure S2. Comparison of automated and manual patch clamp recording of KCNQ2/KCNQ3.** Whole cell current density recorded from CHO-Q3 cells electroporated with wild type KCNQ2 (Q2-WT) using either automated (**A**) or manual (**B**) patch clamp.

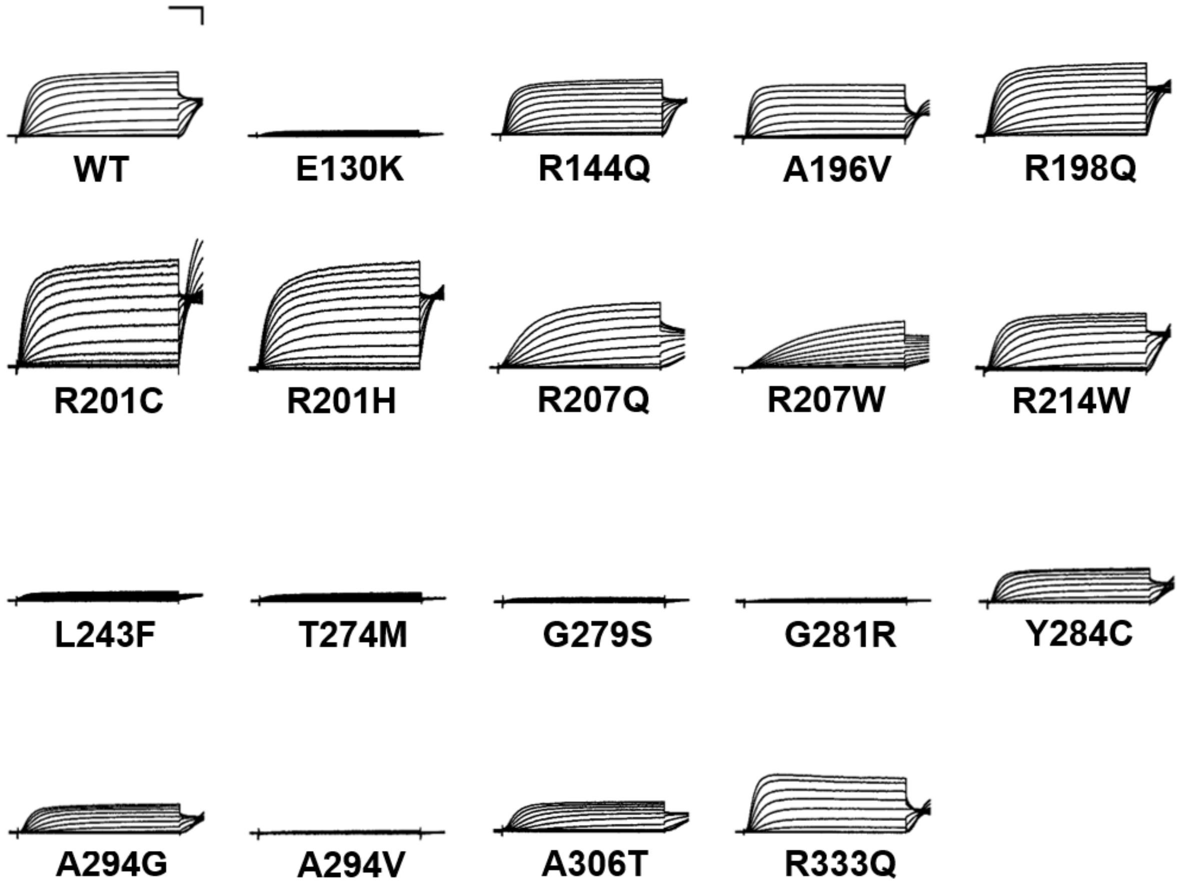


**Figure S3. Whole-cell currents from literature KCNQ2 variants expressed as homozygous channels.** Average XE-991-sensitive whole-cell currents recorded by automated patch clamp from CHO-Q3 cells electroporated with KCNQ2 variants from the literature set and normalized to wild type channel peak current. For variant R201C, whole-cell currents were recorded from CHO-K1 cells co-electroporated with KCNQ3-WT plus KCNQ2-variant. Scale bars are 200 ms (horizontal) and 25% of WT channel current density (vertical).

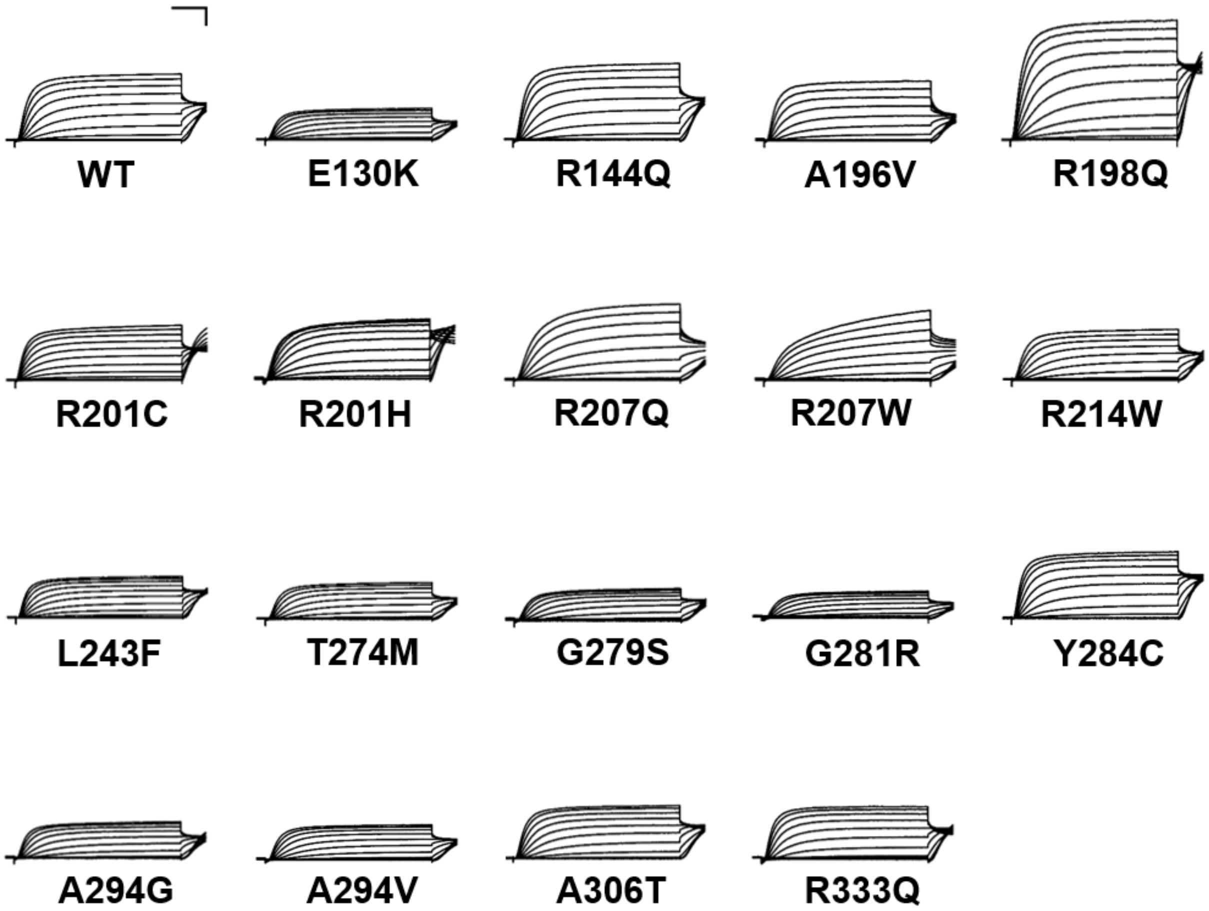


**Figure S4. Whole-cell currents from literature KCNQ2 variants expressed as heterozygous channels.** Average XE-991-sensitive whole-cell currents recorded by automated patch clamp from CHO-Q3 cells co-electroporated with wild type plus variant KCNQ2 cDNA from the literature set and normalized to wild type channel peak current. Scale bars are 200 ms (horizontal) and 25% of WT channel current density (vertical).

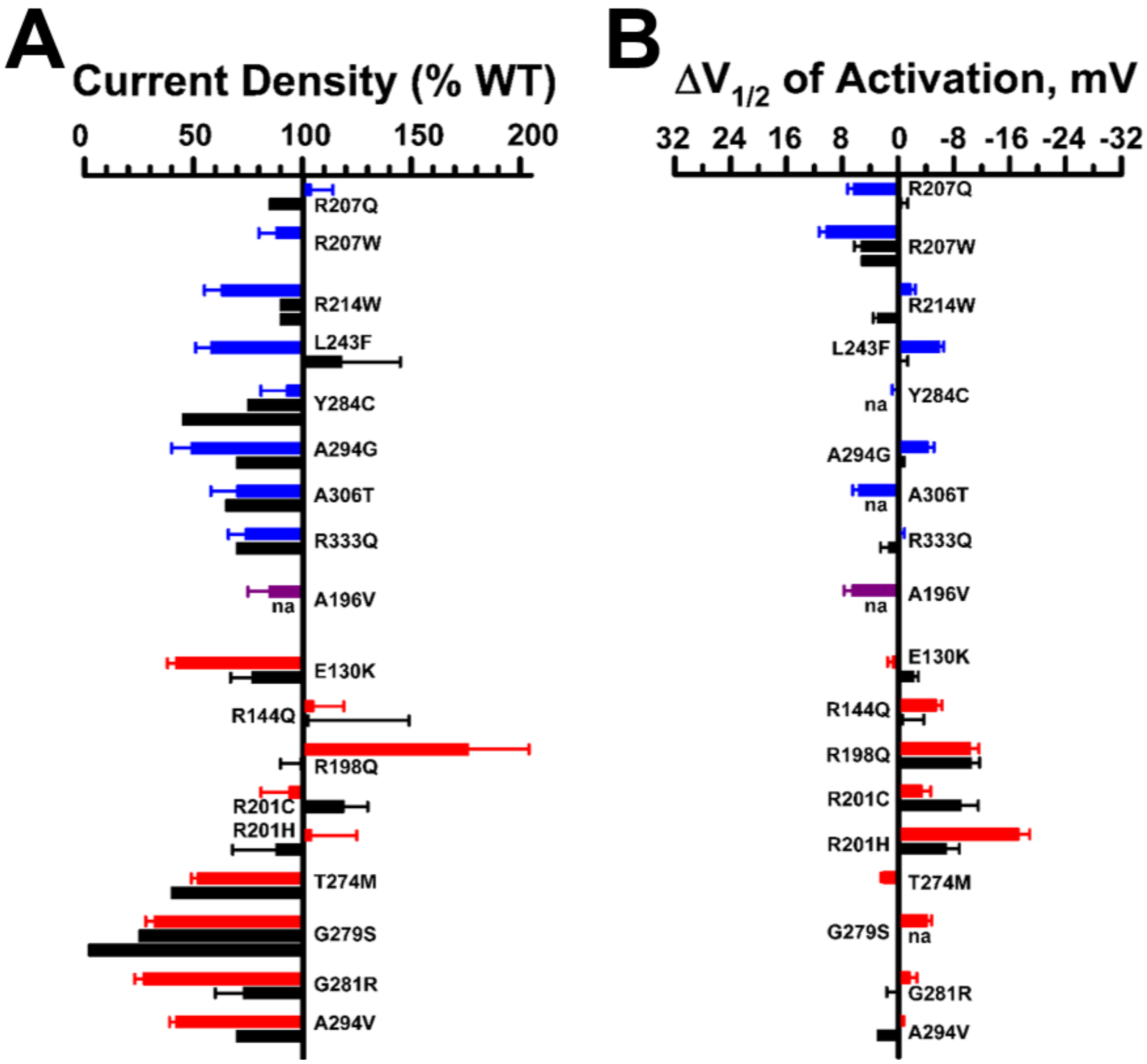


**Figure S5. Manual and automated patch clamp analyses of KCNQ2 variants expressed in the heterozygous state yield similar biophysical properties. A**. Average whole-cell currents recorded at +40 mV from CHO-Q3 cells co-expressing variant + wild type KCNQ2 and normalized to WT channel peak current that was measured in parallel. **B**. Change in current voltage-dependence of activation V½ determined for CHO-Q3 cells co-expressing variant + wild type KCNQ2 relative to WT channel. Black bars indicate literature manual patch clamp data, while blue bars are automated patch clamp results from BFNE-associated variants, red bars represent data from DEE-associated variants, and the purple bar is a BFNE/DEE-associated variant. na = not available in the literature.

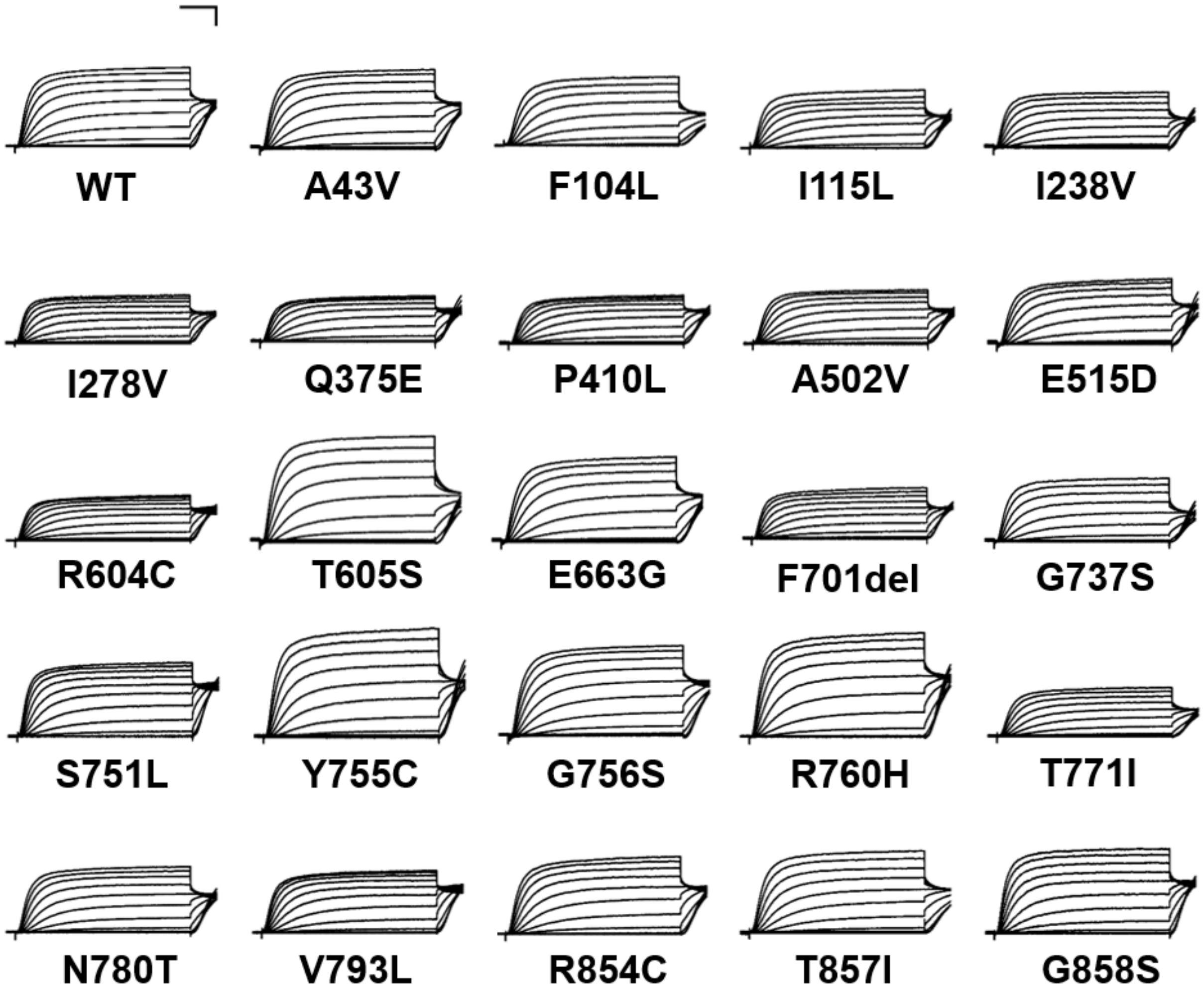


**Figure S6. Average whole-cell currents recorded from CHO-Q3 cells electroporated with population KCNQ2 variants.** Average XE-991-sensitive whole-cell currents recorded by automated patch clamp from CHO-Q3 cells electroporated with rare population KCNQ2 variants and normalized to wild type channel peak current. Scale bars are 200 ms (horizontal) and 25% of WT channel current density (vertical).

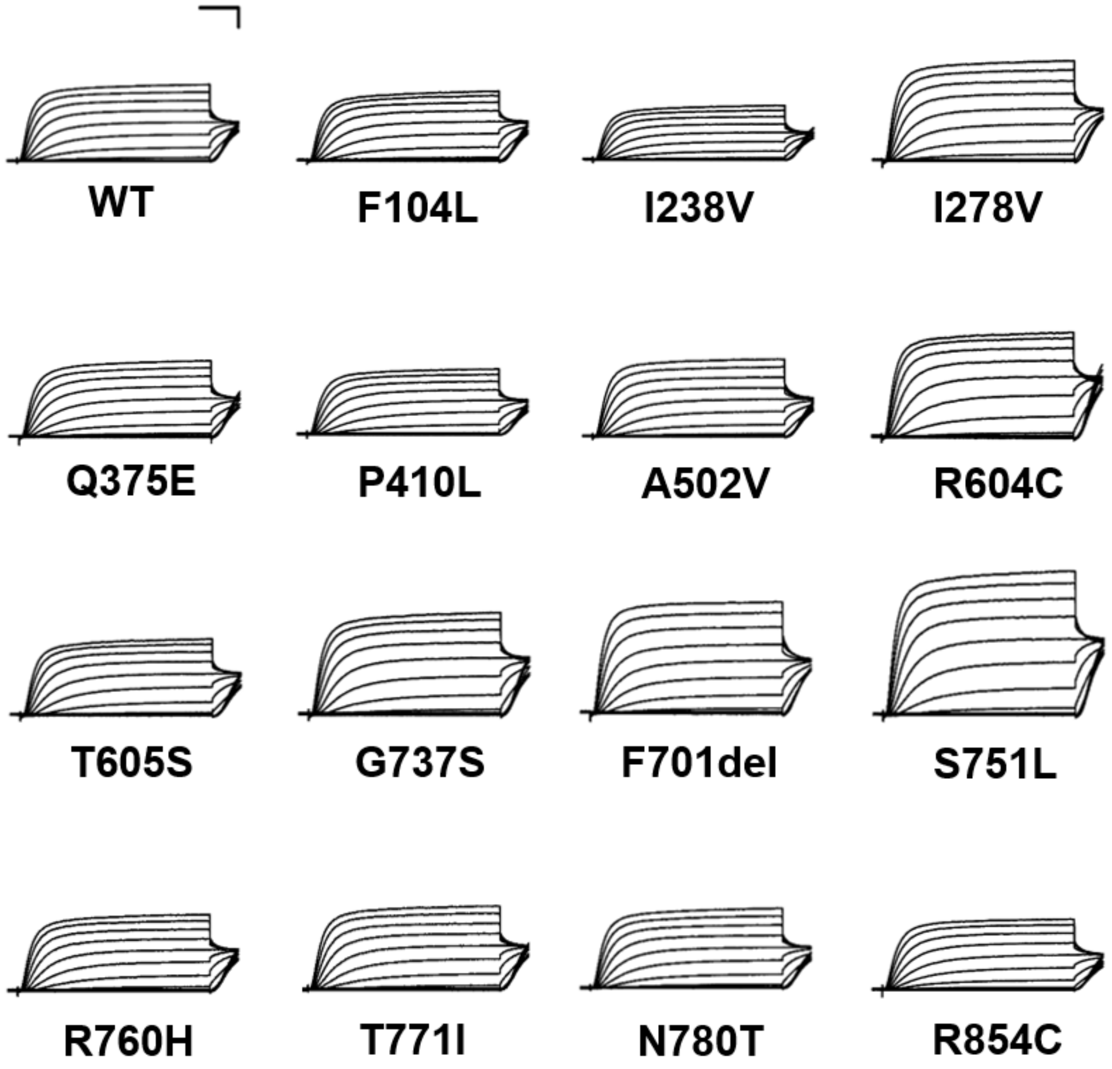


**Figure S7. Average whole-cell currents recorded from CHO-Q3 cells co-electroporated with selected population variants plus wild type KCNQ2.** Average XE-991-sensitive whole-cell currents recorded by automated patch clamp from CHO-Q3 cells co-electroporated with rare population variants plus wild type KCNQ2 and normalized to wild type channel peak current. Scale bars are 200 ms (horizontal) and 25% of WT channel current density (vertical).

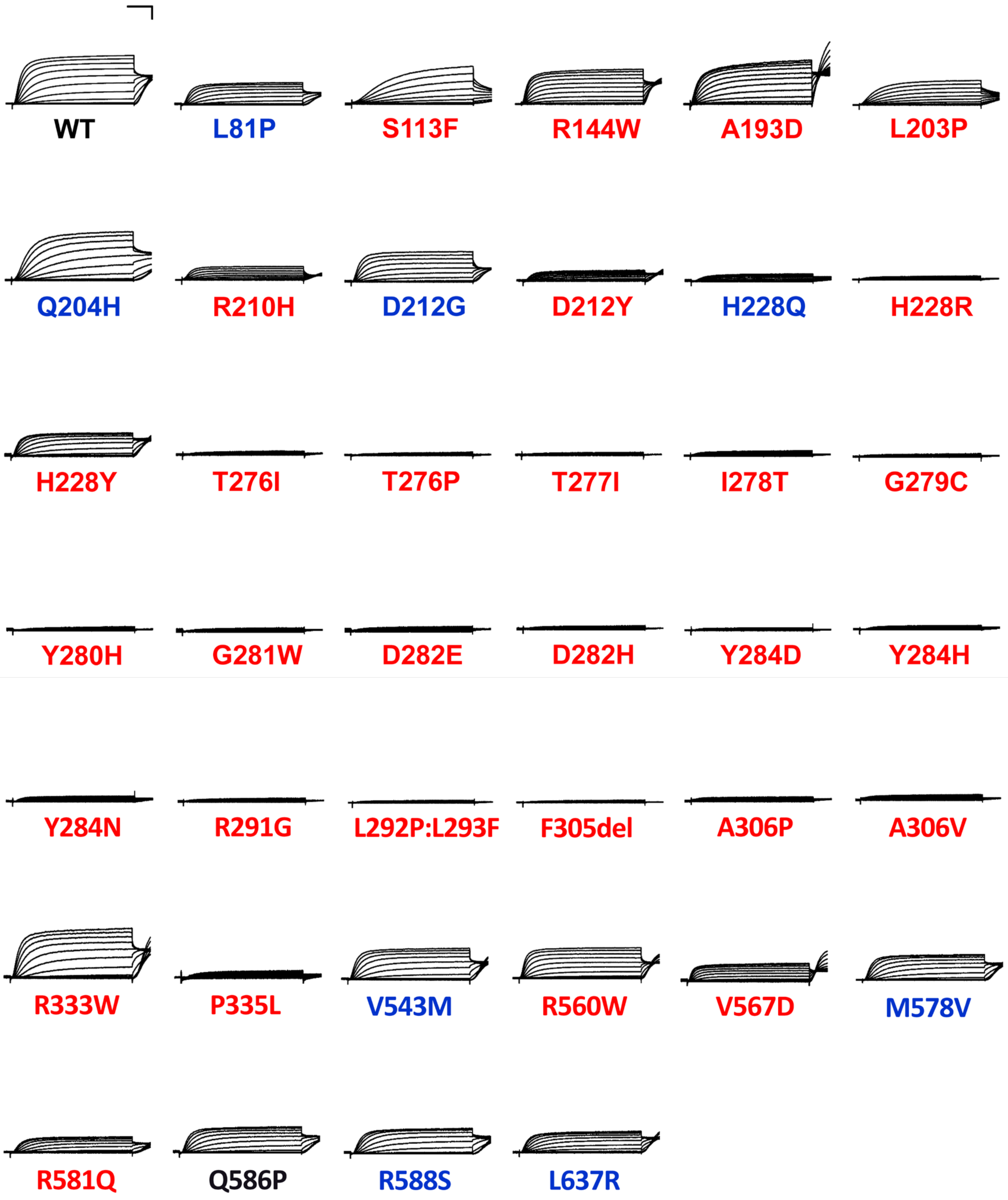


**Figure S8. Whole-cell currents from epilepsy-associated KCNQ2 variants expressed as homozygous channels.** Average XE-991-sensitive whole-cell currents recorded by automated patch clamp from CHO-Q3 cells electroporated with epilepsy-associated KCNQ2 variants and normalized to wild type channel peak current. Variant labels: **Blue** = BFNE-associated; **Red** = DEE-associated; **Black** = unknown phenotype category (Q586P). For A193D and P335L, whole-cell currents were recorded from CHO-K1 cells co-electroporated with KCNQ3-WT plus KCNQ2-variant. Scale bars are 200 ms (horizontal) and 25% of WT channel current density (vertical).

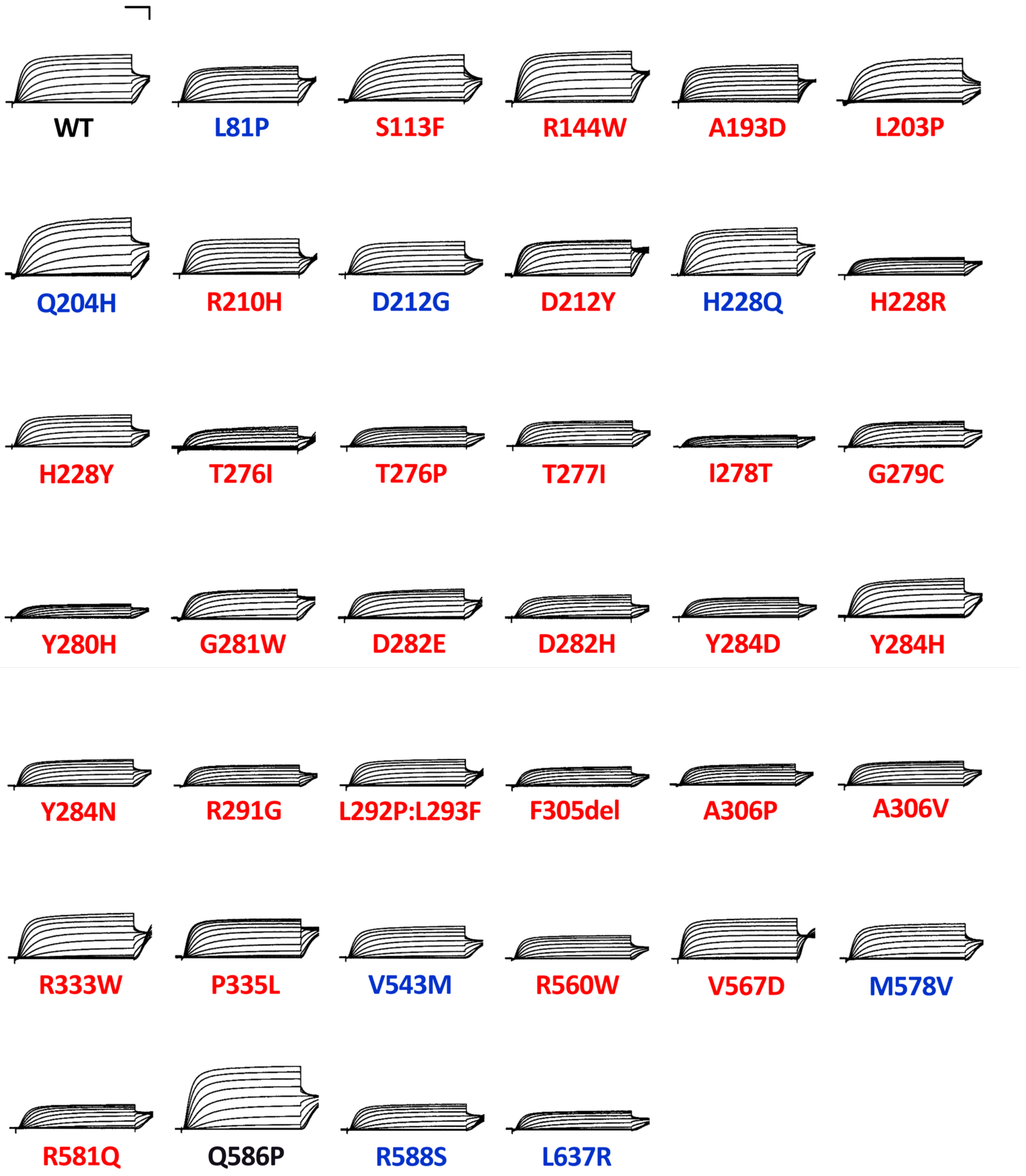


**Figure S9. Average whole-cell currents recorded from CHO-Q3 cells co-electroporated with epilepsy-associated variants plus wild type KCNQ2.** Average XE-991-sensitive whole-cell currents recorded by automated patch clamp from CHO-Q3 cells co-electroporated with epilepsy-associated KCNQ2 variants plus WT KCNQ2 and normalized to wild type channel peak current. Variant labels: **Blue** = BFNE-associated; **Red** = DEE-associated; **Black** = unknown phenotype category (Q586P). Scale bars are 200 ms (horizontal) and 25% of WT channel current density (vertical).

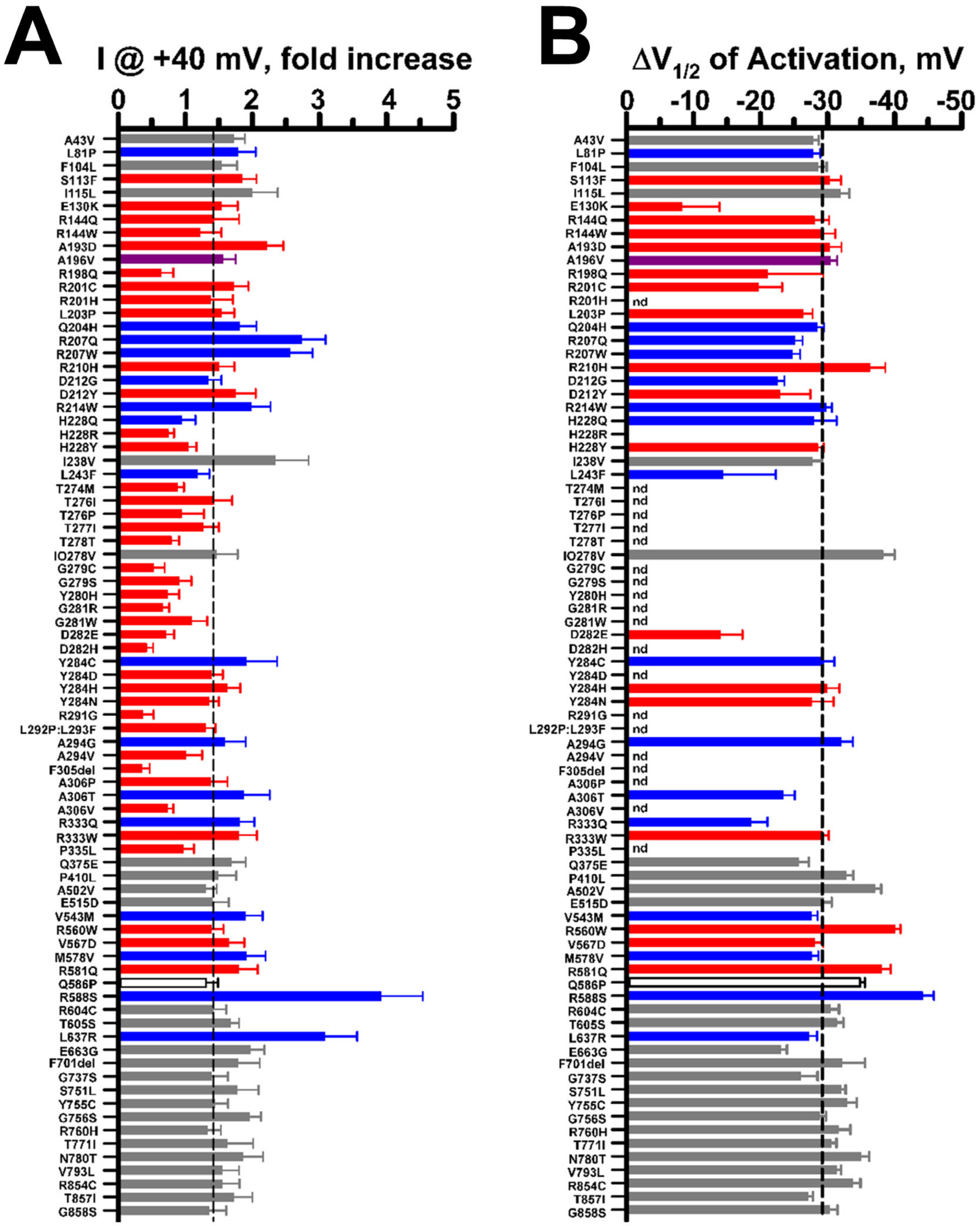


**Figure S10. Retigabine effects on whole-cell currents recorded from KCNQ2 variants expressed in the homozygous state. A**) Ratio of whole-cell currents recorded at +40 mV after exposure to 10 µM retigabine and divided by the current measured under control conditions (n = 15-86). **B**) Change in voltage-dependence of activation V½ determined for whole-cell currents recorded under control conditions and after exposure to 10 µM retigabine (n = 5-74). Dashed lines indicate average effect of retigabine on current amplitude and voltage-dependence of activation V½ in the wild type channel. Variant labels: **Blue** = BFNE-associated; **Red** = DEE-associated.; **Grey** = population variants; unfilled bar = unclear phenotype. For complete list of results, see **Supplemental Table 6**.

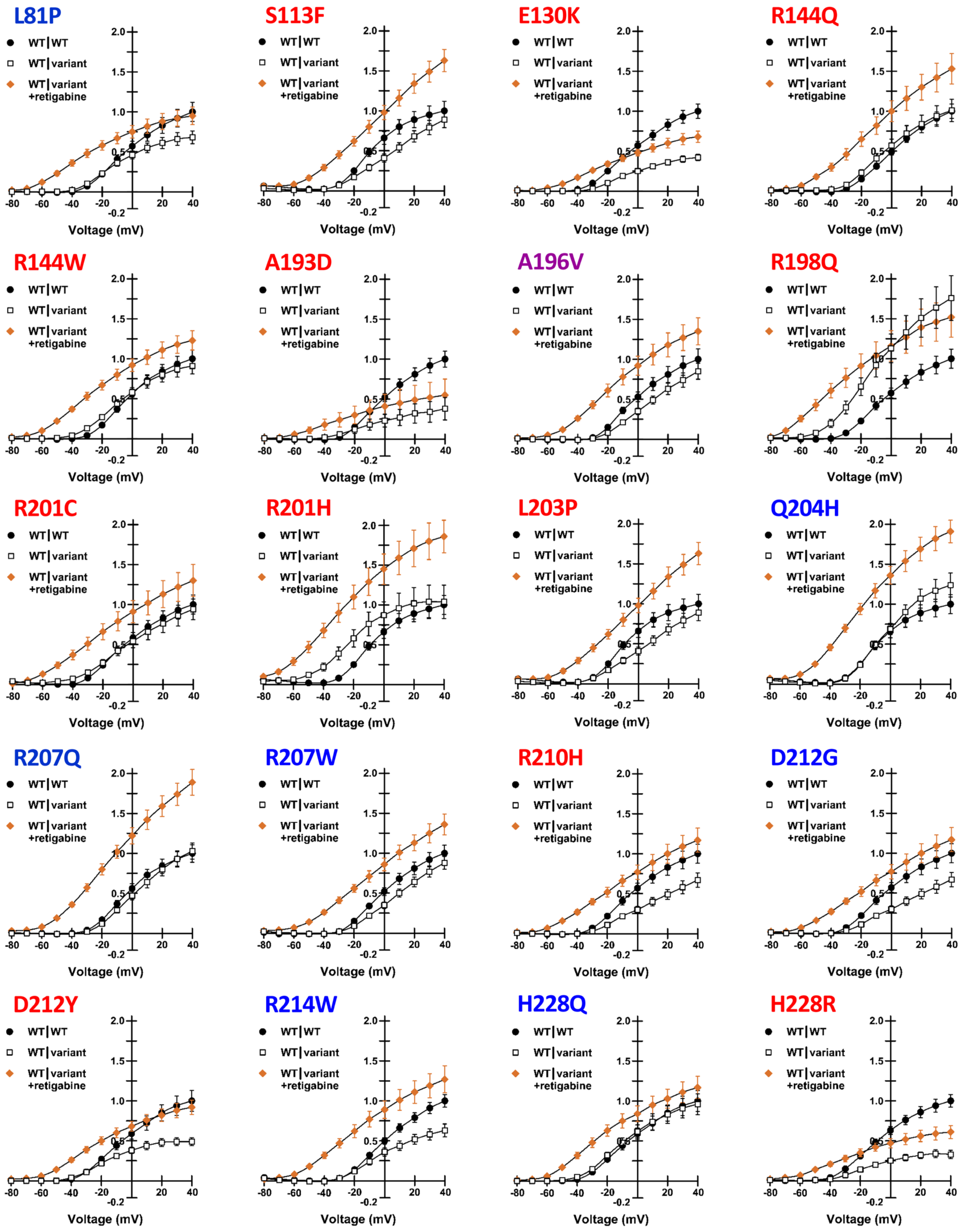


**Figure S11. Retigabine effects on whole-cell currents recorded from epilepsy-associated KCNQ2 variants expressed in the heterozygous state.** Normalized current-voltage relationships for each variant expressed in the heterozygous state recorded in the absence of retigabine (WT|variant, open squares) compared with heterozygous variants recorded in the absence of retigabine (WT|variant +retigabine, orange filled diamonds). Currents were first normalized to cell capacitance, then re-normalized to the peak current for WT channels (WT|WT, filled circles). Variant labels: **Blue** = BFNE-associated; **Red** = DEE-associated; **Purple** = BFNE/DEE; **Black** = unknown phenotype category. Complete data sets are presented in **Table S7**.

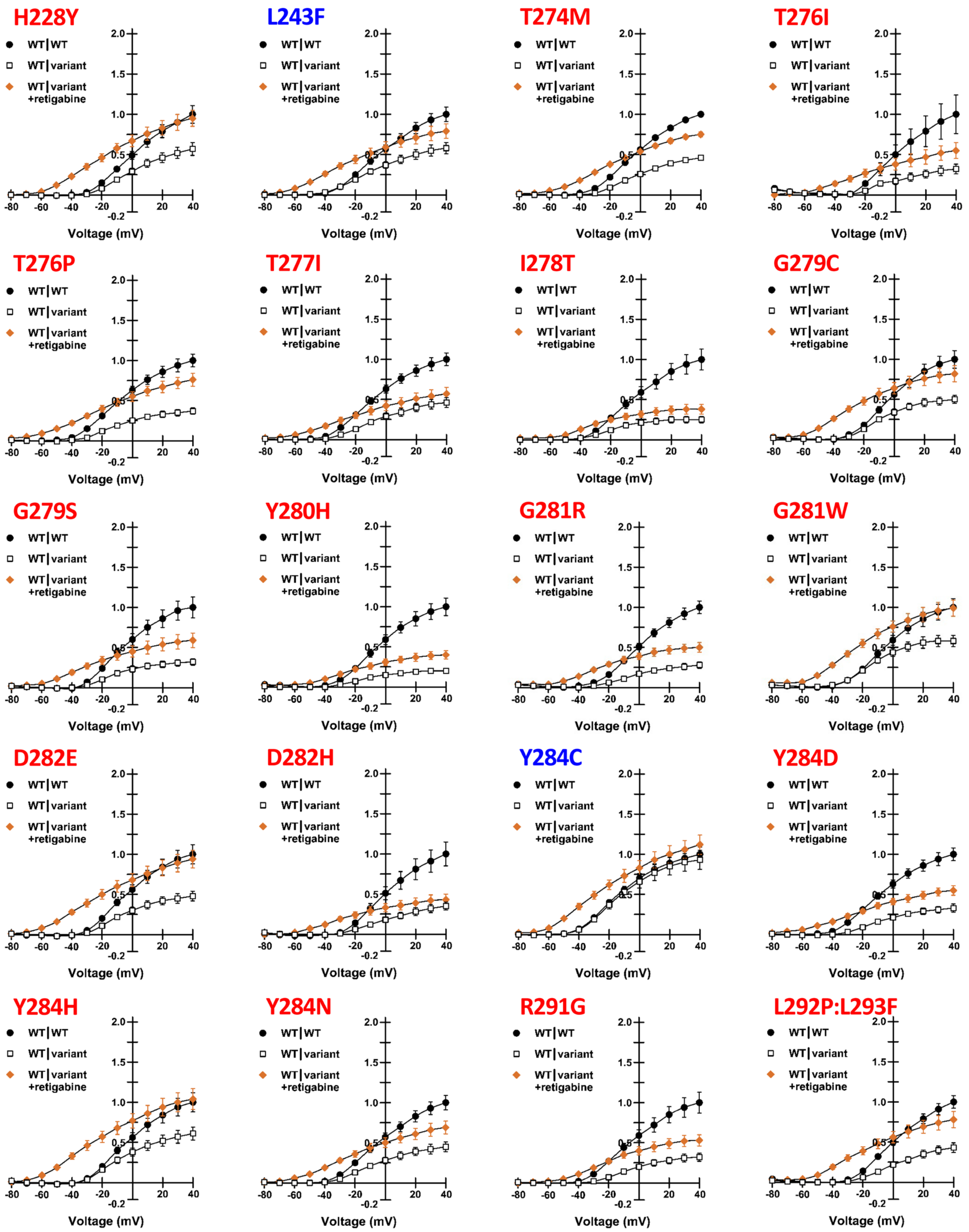


**Figure S11 -** *continued*. **Retigabine effects on whole-cell currents recorded from epilepsy-associated KCNQ2 variants expressed in the heterozygous state.** Normalized current-voltage relationships for each variant expressed in the heterozygous state recorded in the absence of retigabine (WT|variant, open squares) compared with heterozygous variants recorded in the absence of retigabine (WT|variant +retigabine, orange filled diamonds). Currents were first normalized to cell capacitance, then re-normalized to the peak current for WT channels (WT|WT, filled circles). Variant labels: **Blue** = BFNE-associated; **Red** = DEE-associated; **Purple** = BFNE/DEE; **Black** = unknown phenotype category. Complete data sets are presented in **Table S7**.

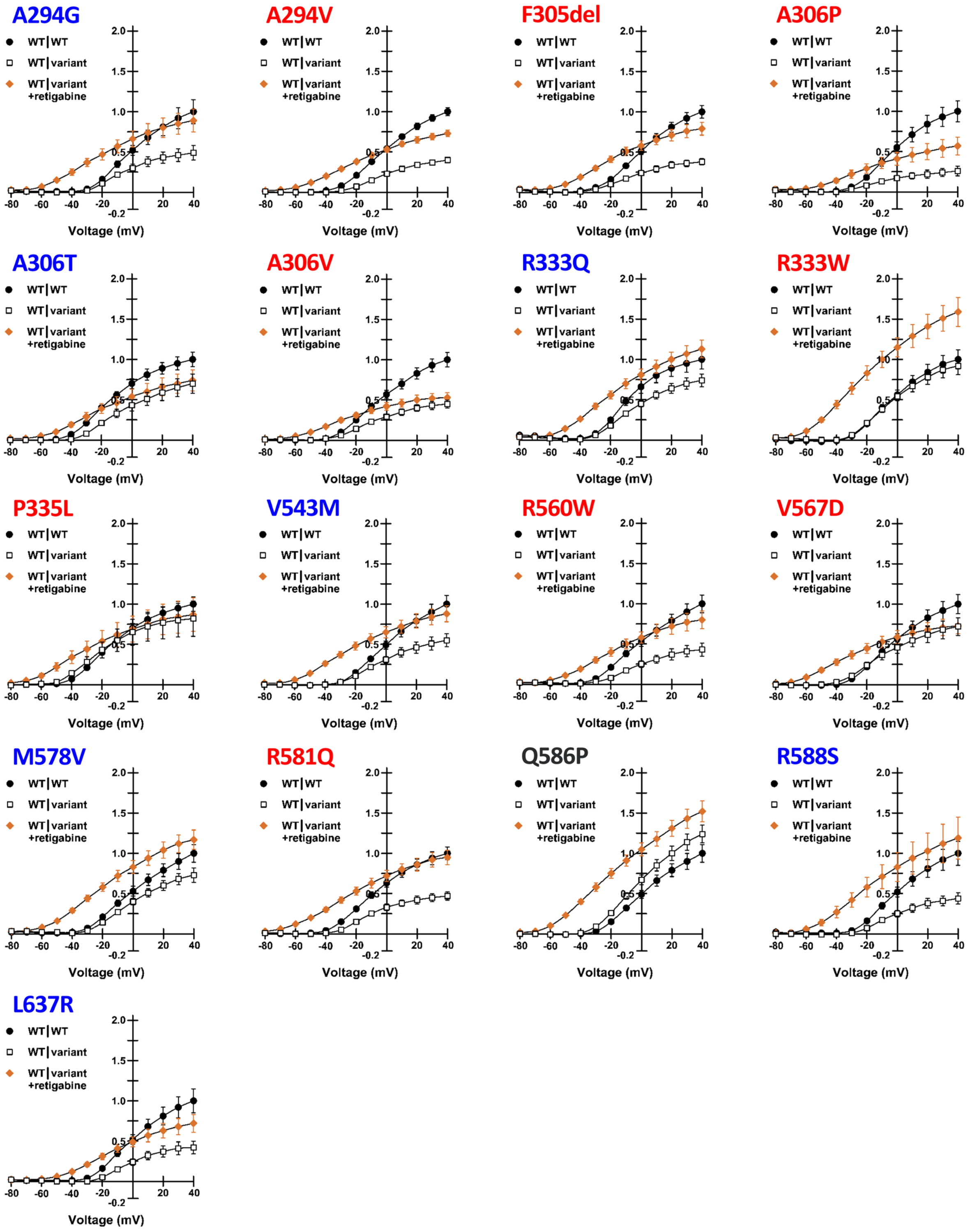


**Figure S11 -** *continued***. Retigabine effects on whole-cell currents recorded from epilepsy-associated KCNQ2 variants expressed in the heterozygous state.** Normalized current-voltage relationships for each variant expressed in the heterozygous state recorded in the absence of retigabine (WT|variant, open squares) compared with heterozygous variants recorded in the absence of retigabine (WT|variant +retigabine, orange filled diamonds). Currents were first normalized to cell capacitance, then re-normalized to the peak current for WT channels (WT|WT, filled circles). Variant labels: **Blue** = BFNE-associated; **Red** = DEE-associated; **Purple** = BFNE/DEE; **Black** = unknown phenotype category. Complete data sets are presented in **Table S7**.

### Table S1 – KCNQ2 variant information

| Nucleotide | Amino Acid | Channel Domain | Phenotype | MAF (gnomAD) | ClinVar | PubMed ID |
|---|---|---|---|---|---|---|
| c.128C>T | p.Ala43Val | N-term | PV | 0.000176 | LB/VUS | |
| c.242T>C | p.Leu81Pro | N-term | BFNE | 0 | N/A | 29215089 |
| c.312C>G | p.Phe104Leu | TMD: S1 | PV | 0.000008 | N/A | |
| c.338C>T | p.Ser113Phe | TMD: S1-S2-Link | DEE | 0 | VUS/LP | 29655203 |
| c.343A>C | p.Ile115Leu | TMD: S1-S2-Link | PV | 0.000016 | N/A | |
| c.388G>A | p.Glu130Lys | TMD: S2 | DEE | 0 | PATH | 27535030 |
| c.431G>A | p.Arg144Gln | TMD: S2-S3-Link | DEE | 0 | PATH/LP | 23934111 |
| c.430C>T | p.Arg144Trp | TMD: S2-S3-Link | DEE | 0 | PATH/LP | 28628100; 28867141 |
| c.578C>A | p.Ala193Asp | TMD: S2-S3-Link | DEE | 0 | PATH | 27602407 |
| c.587C>T | p.Ala196Val | TMD: S4 | DEE | 0 | PATH | 17475800 |
| c.593G>A | p.Arg198Gln | TMD: S4 | DEE | 0 | PATH | 27861786 |
| c.601C>T | p.Arg201Cys | TMD: S4 | DEE | 0 | PATH/VUS | 24107868 |
| c.602G>A | p.Arg201His | TMD: S4 | DEE | 0 | PATH | 23708187 |
| c.608T>C | p.Leu203Pro | TMD: S4 | DEE | 0 | PATH | 26007637 |
| c.612G>T | p.Gln204His | TMD: S4 | BFNE | 0 | LP | 27602407 |
| c.620G>A | p.Arg207Gln | TMD: S4 | DEE | 0 | PATH/LP | 17872363 |
| c.619C>T | p.Arg207Trp | TMD: S4 | BFNE | 0 | PATH | 11572947 |
| c.629G>A | p.Arg210His | TMD: S4 | DEE | 0 | PATH | 24107868 |
| c.635A>G | p.Asp212Gly | TMD: S4 | BFNE | 0 | N/A | 19344764 |
| c.634G>T | p.Asp212Tyr | TMD: S4 | DEE | 0 | PATH | 28817111 |
| c.640C>T | p.Arg214Trp | TMD: S4 | BFNE | 0 | PATH/LP | 11175290; 29056246 |
| c.684C>A | p.His228Gln | TMD: S4-S5-Link | BFNE | 0 | VUS | 14534157 |
| c.683A>G | p.His228Arg | TMD: S4-S5-Link | DEE | 0 | LP | |
| c.682C>T | p.His228Tyr | TMD: S4-S5-Link | BFNE | 0 | Not Provided | 28837158 |
| c.712A>G | p.Ile238Val | TMD: S5 | PV | 0.000008 | VUS | |
| c.727C>T | p.Leu243Phe | TMD: S5 | BFNE | 0 | PATH | 14534157 |
| c.821C>T | p.Thr274Met | TMD: P-loop | DEE | 0 | PATH | 22275249 |
| c.827C>T | p.Thr276Ile | TMD: P-loop | DEE | 0 | PATH | 24463883 |
| c.826A>C | p.Thr276Pro | TMD: P-loop | DEE | 0 | N/A | 29720203 |
| c.830C>T | p.Thr277Ile | TMD: P-loop | DEE | 0 | N/A | 26544041 |
| c.833T>C | p.Ile278Thr | TMD: P-loop | DEE | 0 | LP | 30109124 |
| c.832A>G | p.Ile278Val | TMD: P-loop | PV | 0.000008 | N/A | |
| c.835G>T | p.Gly279Cys | TMD: P-loop | DEE | 0 | PATH | 25959266 |
| c.836G>A | p.Gly279Ser | TMD: P-loop | DEE | 0 | N/A | 27734276 |
| c.838T>C | p.Tyr280His | TMD: P-loop | DEE | 0 | PATH | 27779742 |
| c.841G>A | p.Gly281Arg | TMD: P-loop | DEE | 0 | LP | 24107868 |
| c.841G>T | p.Gly281Trp | TMD: P-loop | DEE | 0 | PATH | 25880994 |
| c.846C>A | p.Asp282Glu | TMD: P-loop | DEE | 0 | N/A | 28133863 |
| c.844G>C | p.Asp282His | TMD: P-loop | DEE | 0 | VUS/LP | 29655203 |
| c.851A>G | p.Tyr284Cys | TMD: P-loop | BFNE | 0 | PATH | 9425895 |
| c.850T>G | p.Tyr284Asp | TMD: P-loop | DEE | 0 | PATH | 27535030 |
| c.850T>C | p.Tyr284His | TMD: P-loop | DEE | 0 | N/A | 29588952 |
| c.850T>A | p.Tyr284Asn | TMD: P-loop | DEE | 0 | N/A | |
| c.871A>G | p.Arg291Gly | TMD: P-loop | DEE | 0 | N/A | 27779742 |
| c.[875T>C:877C>T] | p.Leu292Pro:Leu293Phe | TMD: S6 | DEE | 0 | LP , VUS | |

### Table S1 – *(continued)* KCNQ2 variant information

| Nucleotide | Amino Acid | Channel Domain | Phenotype | MAF (gnomAD) | ClinVar | PubMed ID |
|---|---|---|---|---|---|---|
| c.881C>G | p.Ala294Gly | TMD: S6 | BFNE | 0 | PATH | 17129708 |
| c.881C>T | p.Ala294Val | TMD: S6 | DEE | 0 | PATH | 17129708 |
| c.913_915delTTC | p.Phe305del | TMD: S6 | DEE | 0 | N/A | 28554332; 28728838; 18640800 |
| c.916G>C | p.Ala306Pro | TMD: S6 | DEE | 0 | PATH | 29655203 |
| c.916G>A | p.Ala306Thr | TMD: S6 | DEE | 0 | PATH | 9425895; 26138355 |
| c.917C>T | p.Ala306Val | TMD: S6 | DEE | 0 | PATH | 31152295 |
| c.998G>A | p.Arg333Gln | C-term | BFNE | 0.000004 | PATH/LP | 29215089; 14534157 |
| c.997C>T | p.Arg333Trp | C-term | DEE | 0 | PATH | 16039833 |
| c.1004C>T | p.Pro335Leu | C-term | DEE | 0 | PATH/LP | 28867141 |
| c.1123C>G | p.Gln375Glu | C-term | DEE | 0.000018 | N/A | |
| c.1229C>T | p.Pro410Leu | C-term | PV | 0.000043 | VUS | |
| c.1505C>T | p.Ala502Val | C-term | PV | 0.000036 | VUS | |
| c.1545G>C | p.Glu515Asp | C-term | PV | 0.002517 | B/LB/VUS | 19380078 |
| c.1627G>A | p.Val543Met | C-term | BFNE | 0.000004 | VUS/LP | 28399683 |
| c.1678C>T | p.Arg560Trp | C-term | DEE | 0 | PATH/LP | 22275249 |
| c.1700T>A | p.Val567Asp | C-term | DEE | 0 | LP | 27888506 |
| c.1732A>G | p.Met578Val | C-term | BFNE | 0 | PATH/LP | 25982755 |
| c.1742G>A | p.Arg581Gln | C-term | DEE | 0 | PATH/LP | 27864847 |
| c.1757A>C | p.Gln586Pro | C-term | DEE | 0 | VUS | |
| c.1764A>T | p.Arg588Ser | C-term | BFNE | 0 | PATH | 25982755 |
| c.1810C>T | p.Arg604Cys | C-term | PV | 0.000008 | VUS | |
| c.1814C>G | p.Thr605Ser | C-term | PV | 0.000056 | VUS/LB | |
| c.1910T>G | p.Leu637Arg | C-term | BFNE | 0 | PATH | 25982755 |
| c.1988A>G | p.Glu663Gly | C-term | PV | 0.000047 | N/A | |
| c.2101_2103delTCT | p.Phe701del | C-term | PV | 0.000009 | N/A | |
| c.2209G>A | p.Gly737Ser | C-term | PV | 0.000016 | VUS | |
| c.2252C>T | p.Ser751Leu | C-term | PV | 0.000065 | VUS | |
| c.2264A>G | p.Tyr755Cys | C-term | PV | 0.002953 | B/LB | |
| c.2266G>A | p.Gly756Ser | C-term | PV | 0.000264 | LB | |
| c.2279G>A | p.Arg760His | C-term | PV | 0.000059 | VUS | |
| c.2312C>T | p.Thr771Ile | C-term | PV | 0.000047 | VUS | |
| c.2339A>C | p.Asn780Thr | C-term | PV | 0.609194 | B | |
| c.2377G>C | p.Val793Leu | C-term | PV | 0.000025 | VUS | |
| c.2560C>T | p.Arg854Cys | C-term | PV | 0.000226 | B/LB | |
| c.2570C>T | p.Thr857Ile | C-term | PV | 0.000019 | N/A | |
| c.2572G>A | p.Gly858Ser | C-term | PV | 0.000030 | VUS | |

### Table S2. Sequence of mutagenic primers used to generate KCNQ2 variants.

| Nucleotide change | Amino acid change | Forward Primer | Reverse Primer |
|---|---|---|---|
| c.128C>T | p.Ala43Val | GCTGATCG**T**CGGCTCCGAGGCCCCCAAG | CGGAGCCG**A**CGATCAGCAGCGCCCCGTCC |
| c.242T>C | p.Leu81Pro | GCAGAATTTCC**C**CTACAACGTGCTGGAGCGGCC | TGTAG**G**GGAAATTCTGCAGCTTGCGGTAGAAGG |
| c.312C>G | p.Phe104Leu | CTGGTTTT**G**TCCTGCCTCGTGCTGTCTGTGTTTTC | AGGCAGGA**C**AAAACCAGGAGGAACACGTAGGCGTG |
| c.338C>T | p.Ser113Phe | GTGTTTT**T**CACCATCAAGGAGTATGAGAAGAGCTCG | TTGATGGTG**A**AAAACACAGACAGCACGAGGCAGGAG |
| c.343A>C | p.Ile115Leu | TTCCACC**C**TCAAGGAGTATGAGAAGAGCTCGGAGG | ACTCCTTGA**G**GGTGGAAAACACAGACAGCACGAGG |
| c.388G>A | p.Glu130Lys | CCTG**A**AAATCGTGACTATCGTGGTGTTTGGCGTG | TAGTCACGATTT**T**CAGGATGTAGAGGGCCCCCTC |
| c.430C>T | p.Arg144Trp | GTACTTCGTG**T**GGATCTGGGCCGCAGGCTGC | AGATCC**A**CACGAAGTACTCCACGCCAAACACC |
| c.431G>A | p.Arg144Gln | TACTTCGTGC**A**GATCTGGGCCGCAGGCTGCTG | CAGATC**T**GCACGAAGTACTCCACGCCAAACAC |
| c.578C>A | p.Ala193Asp | CGTCTTTG**A**CACATCTGCGCTCCGGAGCCT | CAGATGTG**T**CAAAGACGTTGCCCTGGGAGCC |
| c.587C>T | p.Ala196Val | GCCACATCTG**T**GCTCCGGAGCCTGCGCTTCC | CGGAGC**A**CAGATGTGGCAAAGACGTTGCCCTG |
| c.593G>A | p.Arg198Gln | TCTGCGCTCC**A**GAGCCTGCGCTTCCTGCAGAT | CGCAGGCTC**T**GGAGCGCAGATGTGGCAAAGAC |
| c.601C>T | p.Arg201Cys | GCCTG**T**GCTTCCTGCAGATTCTGCGGATGATC | CTGCAGGAAGC**A**CAGGCTCCGGAGCGCAGATGT |
| c.602G>A | p.Arg201His | CCTGC**A**CTTCCTGCAGATTCTGCGGATGATCC | TCTGCAGGAAG**T**GCAGGCTCCGGAGCGCAGATG |
| c.608T>C | p.Leu203Pro | CTTCC**C**GCAGATTCTGCGGATGATCCGCATG | GCAGAATCTGC**G**GGAAGCGCAGGCTCCGGAGC |
| c.612G>T | p.Gln204His | CCTGCA**T**ATTCTGCGGATGATCCGCATGGACC | TCCGCAGAAT**A**TGCAGGAAGCGCAGGCTCCGG |
| c.620G>A | p.Arg207Gln | AGATTCTGC**A**GATGATCCGCATGGACCGGCG | GATCATC**T**GCAGAATCTGCAGGAAGCGCAGG |
| c.619C>T | p.Arg207Trp | CAGATTCTG**T**GGATGATCCGCATGGACCGGC | ATCATCC**A**CAGAATCTGCAGGAAGCGCAGGC |
| c.629G>A | p.Arg210His | GCGGATGATCC**A**CATGGACCGGCGGGGAGGC | CCATG**T**GGATCATCCGCAGAATCTGCAGGAAG |
| c.635A>G | p.Asp212Gly | GATCCGCATGG**G**CCGGCGGGGAGGCACCTG | GCCGG**C**CCATGCGGATCATCCGCAGAATCTG |
| c.634G>T | p.Asp212Tyr | TGATCCGCATG**T**ACCGGCGGGGAGGCACCTG | CCGGT**A**CATGCGGATCATCCGCAGAATCTGC |
| c.640C>T | p.Arg214Trp | TGGACCGG**T**GGGGAGGCACCTGGAAGCTGC | GCCTCCCC**A**CCGGTCCATGCGGATCATCCG |
| c.684C>A | p.His228Gln | TATGCCCA**A**AGCAAGGAGCTGGTCACTGCCTG | TCCTTGCT**T**TGGGCATAGACCACAGAGCCCAG |
| c.683A>G | p.His228Arg | CTATGCCC**G**CAGCAAGGAGCTGGTCACTGCC | CCTTGCTG**C**GGGCATAGACCACAGAGCCCAG |
| c.682C>T | p.His228Tyr | CTATGCC**T**ACAGCAAGGAGCTGGTCACTGCC | CCTTGCTGT**A**GGCATAGACCACAGAGCCCAGC |
| c.712A>G | p.Ile238Val | TGGTAC**G**TCGGCTTCCTTTGTCTCATCCTGGC | AGGAAGCCGA**C**GTACCAGGCAGTGACCAGCTCC |
| c.727C>T | p.Leu243Phe | TTCCTTTGT**T**TCATCCTGGCCTCGTTCCTGGTG | AGGATGA**A**ACAAAGGAAGCCGATGTACCAGGCAG |
| c.821C>T | p.Thr274Met | GGTGGGGCCTGATCA**T**GCTGACCACCA | TGGTGGTCAGC**A**TGATCAGGCCCCACC |
| c.827C>T | p.Thr276Ile | CGCTGA**T**CACCATTGGCTACGGGGACAAGTAC | GCCAATGGTG**A**TCAGCGTGATCAGGCCCCACC |
| c.826A>C | p.Thr276Pro | CACGCTG**C**CCACCATTGGCTACGGGGACAAG | CAATGGTGG**G**CAGCGTGATCAGGCCCCACCA |
| c.830C>T | p.Thr277Ile | TGACCA**T**CATTGGCTACGGGGACAAGTACCC | GTAGCCAATG**A**TGGTCAGCGTGATCAGGCCCC |
| c.833T>C | p.Ile278Thr | GACCACCA**C**TGGCTACGGGGACAAGTACCCC | CGTAGCCA**G**TGGTGGTCAGCGTGATCAGGCC |
| c.832A>G | p.Ile278Val | GACCACC**G**TTGGCTACGGGGACAAGTACCCC | CGTAGCCAA**C**GGTGGTCAGCGTGATCAGGCC |
| c.835G>T | p.Gly279Cys | ACCACCATT**T**GCTACGGGGACAAGTACCCCC | CCGTAGC**A**AATGGTGGTCAGCGTGATCAGGCC |
| c.836G>A | p.Gly279Ser | ACCACCATT**A**GCTACGGGGACAAGTACCCCC | CCGTAGC**T**AATGGTGGTCAGCGTGATCAGGCC |
| c.838T>C | p.Tyr280His | ACCATTGGC**C**ACGGGGACAAGTACCCCCAGAC | TCCCCGT**G**GCCAATGGTGGTCAGCGTGATCAG |
| c.841G>A | p.Gly281Arg | ATTGGCTAC**A**GGGACAAGTACCCCCAGACCTG | TTGTCCC**T**GTAGCCAATGGTGGTCAGCGTGATC |
| c.841G>T | p.Gly281Trp | ATTGGCTAC**T**GGGACAAGTACCCCCAGACCTG | TTGTCCC**A**GTAGCCAATGGTGGTCAGCGTGATC |
| c.846C>A | p.Asp282Glu | TACGGGGA**A**AAGTACCCCCAGACCTGGAACGG | GGGTACTT**T**TCCCCGTAGCCAATGGTGGTCAG |
| c.844G>C | p.Asp282His | CTACGGG**C**ACAAGTACCCCCAGACCTGGAACG | GGTACTTGT**G**CCCGTAGCCAATGGTGGTCAGC |
| c.851A>G | p.Tyr284Cys | GGGACAAGT**G**CCCCCAGACCTGGAACGGCAG | CTGGGGG**C**ACTTGTCCCCGTAGCCAATGGTGG |
| c.850T>G | p.Tyr284Asp | GGGGACAAG**G**ACCCCCAGACCTGGAACGGCA | TGGGGGT**C**CTTGTCCCCGTAGCCAATGGTGG |
| c.850T>C | p.Tyr284His | GGGGACAAG**C**ACCCCCAGACCTGGAACGGCA | TGGGGGT**G**CTTGTCCCCGTAGCCAATGGTGG |

### Table S2 - *continued*. Sequence of mutagenic primers used to generate KCNQ2 variants.

| Nucleotide change | Amino acid change | Forward Primer | Reverse Primer |
|---|---|---|---|
| c.850T>A | p.Tyr284Asn | GGGGACAAG**A**ACCCCCAGACCTGGAACGGCA | TGGGGGT**T**CTTGTCCCCGTAGCCAATGGTGG |
| c.871A>G | p.Arg291Gly | ACCTGGAACGGC**G**GGCTCCTTGCGG | CCGCAAGGAGCC**C**GCCGTTCCAGGT |
| c.[875T>C:877C>T] | p.Leu292Pro:Leu293Phe | CAGGC**C**C**T**TTGCGGCAACCTTCACCCTCATCG | TTGCCGCAA**A**G**G**GCCTGCCGTTCCAGGTCTGGG |
| c.881C>T | p.Ala294Val | CTCCTTG**T**GGCAACCTTCACCCTCATCGGTG | AAGGTTGCC**A**CAAGGAGCCTGCCGTTCCAGG |
| c.881C>G | p.Ala294Gly | CTCCTTG**G**GGCAACCTTCACCCTCATCGGTG | AAGGTTGCC**C**CAAGGAGCCTGCCGTTCCAGG |
| c.913_915delTTC | p.Phe305del | TGTCTCCTTC**...**GCGCTGCCTGCAGGCATCTTG | CAGCGC**...**GAAGGAGACACCGATGAGGGTGAAG |
| c.916G>C | p.Ala306Pro | CTCCTTCTTC**C**CGCTGCCTGCAGGCATCTTGG | GCAGCG**G**GAAGAAGGAGACACCGATGAGGGTG |
| c.916G>A | p.Ala306Thr | CTCCTTCTTC**A**CGCTGCCTGCAGGCATCTTGG | GCAGCG**T**GAAGAAGGAGACACCGATGAGGGTG |
| c.917C>T | p.Ala306Val | CCTTCTTCG**T**GCTGCCTGCAGGCATCTTGGG | AGGCAGC**A**CGAAGAAGGAGACACCGATGAGGG |
| c.998G>A | p.Arg333Gln | GAGAAGAGGC**A**GAACCCGGCAGCAGGCCTGAT | GGGTTC**T**GCCTCTTCTCAAAGTGCTTCTGCCTG |
| c.997C>T | p.Arg333Trp | TTGAGAAGAGG**T**GGAACCCGGCAGCAGGCCTG | GTTCC**A**CCTCTTCTCAAAGTGCTTCTGCCTGTG |
| c.1004C>T | p.Pro335Leu | AGGCGGAACC**T**GGCAGCAGGCCTGATCCAGTC | GCTGCC**A**GGTTCCGCCTCTTCTCAAAGTGCTTC |
| c.1123C>G | p.Gln375Glu | TACAGTTCG**G**AAACTCAAACCTACGGGGCCTCC | GTTTGAGTTT**C**CGAACTGTACATGGGCACGGTG |
| c.1229C>T | p.Pro410Leu | AGGACCCCC**T**GCCGGAGCCGTCTCCAAGCC | CTCCGGC**A**GGGGGTCCTTCCTGAAAGCGAG |
| c.1505C>T | p.Ala502Val | GTGCCG**T**GTCACGGCAGAACTCAGAAGCAAGC | CTGCCGTGAC**A**CGGCACCCTTGATGCGGAAAGC |
| c.1545G>C | p.Glu515Asp | CGGAGA**C**GACATTGTGGATGACAAGAGCTGCCC | CCACAATGTC**G**TCTCCGGGGAGGCTTGCTTCTG |
| c.1627G>A | p.Val543Met | CAGAGCC**A**TGTGTGTCATGCGGTTCCTGGTGTC | TGACACACA**T**GGCTCTGATGCTGACTTTGAGGCC |
| c.1678C>T | p.Arg560Trp | GGAGAGCCTG**T**GGCCCTACGACGTGATGGACG | AGGGCC**A**CAGGCTCTCCTTGAACTTCCGCTTG |
| c.1700T>A | p.Val567Asp | ATGGACG**A**CATCGAGCAGTACTCAGCCGGC | TGCTCGATG**T**CGTCCATCACGTCGTAGGGCC |
| c.1732A>G | p.Met578Val | TGGAC**G**TGCTGTCCCGAATTAAGAGCCTGCAG | TCGGGACAGCA**C**GTCCAGGTGGCCGGCTGAGTA |
| c.1742G>A | p.Arg581Gln | TCCC**A**AATTAAGAGCCTGCAGTCCAGAGTGGAC | AGGCTCTTAATT**T**GGGACAGCATGTCCAGGTGGC |
| c.1757A>C | p.Gln586Pro | AGAGCCTGC**C**GTCCAGAGTGGACCAGATCGTGG | TCTGGAC**G**GCAGGCTCTTAATTCGGGACAGCATG |
| c.1764A>T | p.Arg588Ser | TGCAGTCCAG**T**GTGGACCAGATCGTGGGGCGG | GTCCAC**A**CTGGACTGCAGGCTCTTAATTCGGG |
| c.1810C>T | p.Arg604Cys | GGACAAGGAC**T**GCACCAAGGGCCCGGCCGAG | CCTTGGTGC**A**GTCCTTGTCCGTGATCGCTGG |
| c.1814C>G | p.Thr605Ser | AAGGACCGCA**G**CAAGGGCCCGGCCGAGGC | CCCTTG**C**TGCGGTCCTTGTCCGTGATCGC |
| c.1910T>G | p.Leu637Arg | AGAAGC**G**GGACTTCCTGGTGAATATCTACATGCAGC | CAGGAAGTCC**C**GCTTCTTCTCCATGGACAAGACCTGC |
| c.1988A>G | p.Glu663Gly | GGGGCCAAAG**G**GCCGGAGCCGGCGCCGCC | TCCGGC**C**CTTTGGCCCCAAAGTAGGCCTCG |
| c.2101_2103delTCT | p.Phe701del | CCAGAAGAAC**...**TCGGCGCCCCCGGCCGCG | CGCCGA**...**GTTCTTCTGGCCCGTGGAGCTG |
| c.2209G>A | p.Gly737Ser | GGACCAC**A**GCTCCCTGGTGCGCATCCCG | CCAGGGAGC**T**GTGGTCCCCCACGGGGGAG |
| c.2252C>T | p.Ser751L eu | CGAGCGGT**T**GCTGTCCGCCTACGGCGGG | CGGACAGC**A**ACCGCTCGTGGGCAGGCGG |
| c.2264A>G | p.Tyr755Cys | TGTCCGCCT**G**CGGCGGGGGCAACCGCGC | CCCGCCG**C**AGGCGGACAGCGACCGCTCG |
| c.2266G>A | p.Gly756Ser | TCCGCCTAC**A**GCGGGGGCAACCGCGCCA | CCCCCGC**T**GTAGGCGGACAGCGACCGCTC |
| c.2279G>A | p.Arg760His | GGCAACC**A**CGCCAGCATGGAGTTCCTGCG | ATGCTGGCG**T**GGTTGCCCCCGCCGTAGGC |
| c.2312C>T | p.Thr771Ile | CAGGAGGACA**T**CCCGGGCTGCAGGCCCCC | CCCGGG**A**TGTCCTCCTGCCGCAGGAACTC |
| c.2339A>C | p.Asn780Thr | AGGGGA**C**CCTGCGGGACAGCGACACGTC | GTCCCGCAGG**G**TCCCCTCGGGGGGCCTGC |
| c.2377G>C | p.Val793Leu | TCCCGTCC**C**TGGACCACGAGGAGCTGGAGC | GTGGTCCA**G**GGACGGGATGGAGATGGACGTG |
| c.2560C>T | p.Arg854Cys | CCCCGCCA**T**GCTCGGCCACCGGCGAGG | GGCCGAGC**A**TGGCGGGGGCCCGCACGG |
| c.2570C>T | p.Thr857Ile | CGGCCA**T**CGGCGAGGGTCCCTTTGGTGA | ACCCTCGCCG**A**TGGCCGAGCGTGGCGGGG |
| c.2572G>A | p.Gly858Ser | GCCACC**A**GCGAGGGTCCCTTTGGTGACG | GGACCCTCGC**T**GGTGGCCGAGCGTGGCGG |

**Table S3. Manual patch clamp and high throughput functional results.**

To be uploaded at a late date

**Table S4. Functional properties of CHO-Q3 cells electroporated with homozygous variant KCNQ2 cDNA recorded under control conditions.**

To be uploaded at a late date

**Table S5. Functional properties of CHO-Q3 cells co-electroporated with heterozygous variant plus wild type KCNQ2 cDNA recorded under control conditions.**

To be uploaded at a late date

**Table S6 Functional properties of CHO-Q3 cells electroporated with homozygous variant KCNQ2 cDNA recorded following exposure to retigabine.**

To be uploaded at a late date

**Table S7. Functional properties of CHO-Q3 cells co-electroporated with heterozygous variant plus wild type KCNQ2 cDNA recorded following exposure to retigabine.**

To be uploaded at a late date
